# Provision of phosphates by the host supports kleptoplast functionality in photosynthetic sea slugs

**DOI:** 10.1101/2025.10.30.685612

**Authors:** Vesa Havurinne, Heta Mattila, Carolina Esteves, Paulo Cartaxana, Sónia Cruz

## Abstract

Phosphorus and nitrogen are two fundamental macronutrients needed for photosynthesis, and their balance in photosymbiotic relationships, like those between corals and their zooxanthellae, is known to be highly important for maintaining a healthy partnership between the host and the symbiont. Kleptoplasty is a special case of algae-herbivore interaction, where the host animal or protist sequesters the chloroplasts from their prey algae as kleptoplasts inside their own cells. In some kleptoplastic sea slugs, the kleptoplasts can remain functional for months without the help from the algal nucleus, and the fundamental mechanisms in place to achieve this are still largely unsolved. Unlike in purely symbiotic systems like corals, the importance of nutrient balance to kleptoplast longevity has not been addressed in kleptoplastic sea slugs. We studied the phosphate and nitrogen exchange between the sea slug *Elysia timida* and its *Acetabularia acetabulum* derived kleptoplasts in various nutrient depletion and replenishment experiments. Our results show that sea slugs consumed phosphate from their surrounding medium and were able to recover electron transfer rates of kleptoplasts originating from phosphate depleted algae, suggesting that the sea slug provided the acquired organelles with phosphate to the extent that they increased the photosynthetic capacity remarkably well. Additional available phosphate in the water also increased the overall longevity of the kleptoplasts. The sea slugs were not able to recover the photosynthesis of nitrogen-depleted kleptoplasts, and additional dissolved nitrate and ammonium were not beneficial for kleptoplast longevity. Our results suggest that the phosphate-nitrogen balance inside the sea slug cells is mainly limited by phosphate, the addition of which can help the kleptoplasts maintain photosynthesis, perhaps by allowing the use of the glutamine synthetase (GS) and glutamate synthase (GOGAT) nitrogen assimilation pathway to function as an efficient alternative electron sink.

## Introduction

Photosymbiotic relationships between a heterotrophic host and a photosynthetic symbiont are often mutualistic. The host organisms, like corals, anemones, fungi or even salamanders, provide the symbiont alga or cyanobacterium with protection, nutrients and a source of carbon, and the symbiont makes use of the provisions to maintain photosynthesis (Kerney et al. 2011; Davy et al. 2012; Grimm et al. 2021). This allows the normally heterotrophic host to tap almost directly into Earth’s primary energy source: the Sun. Kleptoplasty, where predatory hosts, like certain sacoglossan sea slugs or protists of the SAR (stramenopiles, alveolates, and rhizarians) supergroup, consume their prey algae but retain the algal chloroplasts (Waugh & Clark, 1986; Rumpho et al. 2011; Christa, 2023), is sometimes referred to as a special case of photosymbiosis. The chloroplasts remain inside the host cell as photosynthetically active stolen organelles, being renamed as kleptoplasts, from days to months or even years (Green et al. 2000; Park et al. 2008; Sellers et al. 2014). The symbiotic nature of kleptoplasty does not extend to mutualism, as it is difficult to find benefits to the algal prey in this relationship (Waugh & Clark, 1986). The predatory hosts, on the other hand, have been shown to gain access to sugars and nitrogen originating from photosynthetic carbon fixation and nitrogen assimilation reactions of their kleptoplasts (Trench et al. 1972; Christa et al. 2014; Cruz et al. 2020; Cartaxana et al. 2021; Karnkowska et al. 2023; Lopes et al. 2025; Rao et al. 2025).

One of the main mysteries of kleptoplasty is the fact that kleptoplasts can remain operational inside a foreign host with limited, like in the case of some kleptoplastic protists (Johnson et al. 2023; Rao et al. 2025), or no access to the original algal nucleus and its gene products (Rumpho et al. 2000). This contradicts our knowledge on photosynthesis; the photosynthetic machinery, photosystem II (PSII) in particular, should be in constant need of nucleus-assisted regulation and repair to dampen the inevitable (oxidative) damage associated with the strong redox reactions driving photosynthetic electron transfer (Terashima et al. 1994; Tyystjärvi & Aro, 1996; Komenda et al. 2024; Tiwari et al. 2024). In kleptoplastic sea slugs, attempts to explain this contradiction include, for example, a high genetic autonomy of the kleptoplasts derived from their specific prey algae (de Vries et al. 2013; Havurinne et al. 2021), the usage or even enhancement of the native photoprotective mechanisms of the chloroplasts by the sea slugs (Christa et al. 2018; Cartaxana et al. 2019; Havurinne & Tyystjärvi, 2020), or by photoprotective behavior of the animals (Cartaxana et al. 2019; Burgués Palau et al. 2024).

The widely publicized suggestion that the sea slugs might have gained important genes for kleptoplast maintenance via horizontal gene transfer from their prey algae (Rumpho et al. 2008) has been shown to be unsupported by later genomic and transcriptomic analyses of the sea slugs (Bhattacharya et al. 2013; Maeda et al. 2021; Eastman et al. 2023). However, it is still possible that the sea slugs supply the kleptoplasts with advantageous proteins, albeit not of algal origin; dual-targeting of nucleus-encoded proteins to both mitochondria and chloroplasts is known to take place in algae, and a similar mechanism could exist in kleptoplastic animals. This idea was put forward decades ago (Rumpho et al. 2000) and recently revisited by considering also the role of protein targeting sequence charge as a determining factor in protein localization (Melo Clavijo et al. 2023). A recent landmark study showed that the kleptoplasts reside in specific phagosome-derived organelles, kleptosomes, inside the sea slug cells (Allard et al. 2025). In the same study, the overwhelming majority of proteins in the combined proteome of the kleptosomes and the kleptoplasts were shown to be of animal origin, but it is still unclear if the animal proteins are inside the kleptoplasts, or just in the surrounding kleptosome. Importantly, if the kleptoplasts do contain animal proteins, those proteins would have to be targeted also through the kleptosome membrane. Another intriguing possibility is that the animals could supply the kleptoplasts with specific complex polyketides, produced only by kleptoplastic sea slugs, that could function as antioxidants protecting the kleptoplasts and the animals themselves from oxidative stress (Torres et al. 2020).

One fundamental aspect of interaction between the host and its kleptoplasts that must take place for photosynthesis to function for months has been left with little attention when it comes to kleptoplast maintenance: nutrient exchange. A certain amount of give-and-take of nutrients must exist between the host and the kleptoplast. The fact that the kleptoplasts fix carbon via photosynthesis is telling of such interactions; as a bare minimum, the hosts must provide the kleptoplasts with an access to inorganic carbon (Trench et al. 1972). Carbon fixation in the Calvin-Benson-Bassham (CBB) cycle, however, also uses phosphate in the form of ATP. This implies that also either ATP itself, or inorganic phosphate needed for ATP production via photophosphorylation, must be transported into the kleptoplasts. Furthermore, ATP is a ubiquitously required cellular energy carrier, and protein phosphorylation is a major constituent in multiple regulation and acclimation mechanisms of the chloroplast, like PSII repair (Longoni & Goldschmidt-Clermont, 2021; Komenda et al. 2024). ATP is also exchanged between mitochondria and chloroplasts in algae, making it an important molecule in the energetic coupling of these organelles (Burlacot et al. 2019). Besides phosphorus, another important macronutrient for maintaining photosynthesis is nitrogen. In chloroplasts, nitrogen is essential for the synthesis of various biomolecules, like amino acids and proteins, and it is also a major component of chlorophyll (Chl) (Bryant et al. 2020; Zhao et al. 2020). The sea slugs *Elysia chlorotica* and *Elysia crispata* have been shown to carry out protein synthesis in their kleptoplasts, which hints that some amount of nitrogen available to the sea slug is used by the kleptoplasts (Mujer et al. 1996; Allard et al. 2025).

Despite their importance, the roles of phosphate and nitrogen in kleptoplast maintenance have not been studied in detail. Here, we used the kleptoplastic sea slug *Elysia timida* and its prey green macroalga *Acetabularia acetabulum* in various nutrient depletion and replenishment experiments to inspect if, and to what extent, the host animal provides the kleptoplasts with these nutrients either from its own reserves or by mediating the delivery of exogenously applied nutrients. Our results show that available excess phosphate lengthens the period of photosynthetic activity of the kleptoplasts and that the sea slugs can recover the photosynthetic activity of phosphate-depleted kleptoplasts using only their own phosphate reserves. Contrary to phosphates, exogenously provided nitrate or ammonium did not benefit the kleptoplasts. Furthermore, the sea slugs were not able to recover the photosynthetic activity of nitrogen-depleted kleptoplasts, suggesting that even if there is a benefit to be had from providing the kleptoplasts with nitrogen, there is a threshold of dysfunction after which the kleptoplasts in *E. timida* become irreparable.

## Materials and methods

### Organisms and culture conditions

The sea slug *E. timida* was collected originally from the Mediterranean Sea (Elba, Italy) (Havurinne & Tyystjärvi, 2020) and continuously maintained in the lab in aerated transparent plastic tanks (2-5 L) filled with artificial sea water (ASW; Red Sea salt, Red Sea Europe, Verneuil d’Avre et d’Iton, France) with a salinity of 35-36 parts per thousand (PPT). The alga *A. acetabulum* was also maintained in transparent plastic tanks of a similar volume, but without aeration and in ASW (PPT 35-36) supplemented with f/2 nutrients (Guillard, 1975). Both the sea slugs and the algae were kept in a 12/12h day/night cycle under PPFD (photosynthetic photon flux density) 30-50 µmol m^-2^s^-1^ at 20-22 °C. The plastic tanks for the sea slugs and the algae were always closed with a transparent lid. The lids of the *E. timida* tanks had small holes in them to allow entrance for the silicon tubes delivering air to the spargers inside. The sea slugs were fed once every one to two weeks, and only healthy, green adult sea slugs (5-10mm in length) were used in the experiments unless mentioned otherwise.

### *Elysia timida* starvation and re-feeding

Altogether five separate starvation experiments involving *E. timida* were done in this study. The first three experiments involved fed sea slugs taken directly from their growth conditions, whereas the two last ones involved sea slugs that were bleached of their kleptoplasts with strong light and then re-fed with specifically treated algae. After the initial feeding, the sea slugs were kept away from their food source, i.e. let to starve. In all starvation experiments, the animals were placed in new tanks each week and given fresh ASW with their respective nutrients (see below), and they all took place in the growth conditions of the sea slugs. Photosynthetic activity was measured in the sea slugs at different time points as indicated in the figures and text. Once the PSII activity parameter ΔF/F_M_’ (see “Photosynthesis measurements” below for the definition) of an individual sea slug reached zero during the starvation, it was treated as zero in statistical analyses for the rest of the starvation experiment even if the individual perished later, maintaining a constant number of replicates for the entirety of the experiment for all treatment groups.

In the first starvation experiment, 40 sea slugs were distributed to two tanks (20 individuals per tank), each containing 2L of ASW. One of the tanks was spiked with 36 μM NaH₂PO₄·H₂O (+P group), the same concentration as in the f/2 medium of *A. acetabulum*, while the other one contained plain ASW (control). The addition of NaH₂PO₄·H₂O lowered the pH of the ASW from 8.08 (SD±0.01, n=3) to 8.04 (SD±0.01, n=3); due to the small difference, the pH was not controlled in either treatment. In the second starvation experiment, the setup was identical to the first one, but instead of NaH₂PO₄·H₂O, one of the tanks was spiked with 882 μM NaNO_3_ (+NaNO_3_ group), again using the same concentration as in f/2 medium, and the other tank was filled with ASW (control). The third starvation experiment also involved the addition of a nitrogen source to the treatment group, but instead of NaNO_3_, NH_4_Cl was used (+NH_4_Cl group). The final concentration of NH_4_Cl was 20 µM, chosen based on the previous nitrogen assimilation experiments with *E. timida* (Cartaxana et al. 2021).

The bleaching of *E. timida* of their old kleptoplasts for the fourth and fifth starvation experiments was done by placing sea slugs under strong light (PPFD ∼ 1500-2000 µmol m^-2^s^-1^; 12/12h day/ night) for 10-14 days until the animals were estimated by eye to contain little to no green coloration, as previously described (Havurinne et al. 2025). The bleached *E. timida* individuals were fed with *A. acetabulum* depleted of phosphorus or nitrogen (or non-depleted algae as a control; see “*Acetabularia acetabulum* nutrient depletion and repletion” below) for seven days in nets placed inside a water-circulating 150 L life support system (LSS) filled with ASW. The ASW in the LSS was checked weekly for the absence of nitrogen and phosphate sources using commercial aquarium nutrient test kits (Tropic Marin, Hünenberg, Switzerland). The temperature in the LSS was 20 °C and the light PPFD was 50 µmol m^-2^s^-1^, operated at a 12/12h day/night cycle. Photosynthesis was measured from the animals after one, three and seven days of feeding on their respective algae. The re-fed *E. timida* individuals were placed inside 5 L plastic boxes to starve in ASW without additional phosphate (-P group) or in ASW spiked with 36 μM NaH₂PO₄·H₂O (+P group). This starvation experiment was done twice due to the limited amount of sea slugs available for the first experiment.

### *Acetabularia acetabulum* nutrient depletion and repletion

To deplete *A. acetabulum* of specific nutrients, the algae were placed in transparent 1L plastic containers filled initially with 200mL of ASW containing either all of the f/2 nutrients (control), all nutrients except NaNO_3_ (-N), or all nutrients except NaH₂PO₄·H₂O (-P). The bottom of each tank was almost entirely covered by the algae at different stages of their life cycle, and all tanks contained approximately similar amounts of algae. To minimize the high risk of contamination during the long nutrient depletion experiments, the algae were never taken out of their tanks, and the ASW+f/2 media were never changed completely. Instead, 50mL of their respective sterile media was added to each tank once every week. Photosynthesis was estimated from the algae on a weekly basis to inspect them for robust defects in their photosynthetic machinery (see “Photosynthesis measurements” below). Once the specific depletion treatments were deemed to have caused clear hindrances to the photosynthetic electron transfer of the algae, most of them were fed to the bleached sea slugs (see “*Elysia timida* starvation and re-feeding” above), but some of the algae were placed in Petri dishes filled with ASW+f/2 with all the nutrients for three days in the growth conditions to inspect if the defects in photosynthesis were recoverable. The capability of nitrogen-depleted algae to recover using nitrogen in the form of NH_4_Cl was tested by substituting the nitrogen source of f/2, NaNO_3_, with 20 µM NH_4_Cl in otherwise complete ASW+f/2 medium.

### Pigment analysis

The fresh weight of *A. acetabulum* was estimated by gently drying the algae with tissue paper and weighing the dried algal material on an analytical scale. The algae were then placed in a Dounce tissue grinder with 80% acetone and homogenized. The pigment extract was centrifuged (10,000x*g* for five min) and the supernatant was collected. Total Chl was quantified spectrophotometrically immediately afterwards using the formulae and extinction coefficients described by Wellburn (1994).

### Photosynthesis measurements

Photosynthesis was estimated from all sea slugs and algae fluorometrically by measuring rapid light curves (RLCs) with an Imaging-PAM (Mini version; Heinz Walz GmbH, Effeltrich, Germany), unless mentioned otherwise. Each light step of the RLCs lasted 60 s and a saturating light pulse was fired at the end of each step to calculate relative electron transfer rate (rETR) of PSII, defined as 0.42 x ((F_M_’-F)/F_M_’) x PPFD, where 0.42 is a plant-based estimation of the fraction of incident photons absorbed by PSII, and F_M_’ and F are the maximum and transient fluorescence of samples under illumination. RLCs were only analyzed up to the PPFD where all biological replicates of the same treatment exhibited clear, non-zero rETR values.

The samples for the RLCs were taken from their growth/experimental conditions between the hours 10:00-16:00 (growth lights on between 8:00-20:00) without dark acclimation, and the effective PSII photochemical efficiency, ΔF/F_M_’ (i.e. (F_M_’-F)/F_M_’)) at the start of the RLCs was used as a general indicator of PSII photochemical activity. All RLCs were modeled to obtain the maximum relative electron transfer of PSII (rETR_MAX_) for additional comparisons (Eilers & Peeters, 1988). The modeling was done using the Solver plug-in of MS Excel v.16.0 (MS Office 365; Microsoft Corporation, Redmond, WA, USA).

With the algae, the RLCs were measured by placing the containers or Petri dishes without their lids under the measuring head of the Imaging-PAM and adjusting the distance so that the majority of the algae were in focus. The saturating light pulse intensity setting was 10 (maximum allowed) and the duration was 600 ms. The measuring beam intensity setting varied between 1-3 depending on the algae, and the measuring light frequency was always set to 1 Hz. For the RLC analysis, areas of interest from four individual algae were selected for each treatment, and each individual alga was treated as a biological replicate.

Two different methods were used to immobilize sea slugs for the RLC measurements. The first method relied on encasing the sea slugs in 0.2% agar (made in ASW) as described earlier (Morelli et al. 2024), but due to the likely stress caused to the animals by the agar confinement, the same individuals could not be used in any further measurements, decreasing the number of animals in later time points of the starvation experiments. During the course of the study, a less stressful immobilization method was developed, where the sea slugs were pipetted on a matte black cloth and the residual ASW was sucked out from below by a tissue paper, rendering the animals immobile and for the duration of the measurements, while still sufficiently moist to not cause them excessive harm. Due to its non-invasive nature, the animals immobilized with this method were released back into their respective starvation or treatment conditions and used for measurements in other time points of the same experiments; the black cloth method was used in most of the measurements. The agar method was used in some of the measurements in the *E. timida* re-feeding experiment (see “*Elysia timida* starvation and re-feeding” above). The immobilization method itself did not affect the RLC measurements in *E. timida,* as both methods resulted in very similar RLCs when tested (Supplementary Fig. S1).

The light settings of Imaging-PAM used for the sea slugs were identical to the ones used for the algae, except that the measuring light intensity settings varied between 3-6. The areas of interest selected from one individual sea slug were treated as a representation of one biological replicate for the RLC analyses.

### Phosphate uptake by *Elysia timida*

Phosphate uptake by *E. timida* from the surrounding ASW in normal day/night cycle or in continuous darkness was inspected using a commercial aquarium phosphate detection kit (Tropic Marin PO_4_ Test; Tropic Marin). In total, 120 *E. timida* individuals were collected and kept in a beaker with ASW for 3 h. The sea slugs were subsequently divided into six flasks (20 individuals per flask) containing 50 mL ASW supplemented with 36 μM NaH₂PO₄·H₂O. Three flasks were put to regular growth conditions and three flasks were covered with aluminum foil to keep the sea slugs in the dark. As a control, the ASW where the 120 sea slugs had been in for 3 h was also divided into three additional flasks, 50 mL in each, and also supplemented with 36 μM NaH₂PO₄·H₂O. No sea slugs were placed in the control flasks, kept in growth conditions. All flasks with and without sea slugs were aerated through a pipette tip attached to a silicon hose and an air pump for the whole duration of the experiment. The flasks were sealed with parafilm, but the silicon hose pass-through did leave a small opening for water evaporation.

Five mL of ASW was taken from each flask after four and seven days to estimate the phosphate concentrations according to the Tropic Marin PO_4_ Test instructions. Instead of relying on the comparator provided by the test kit, photographs were taken from the samples after the final incubation step of the test protocol to compare the phosphate concentrations of the samples between each other and to a dilution series made at the beginning of the experiment from the ASW containing 36 μM NaH₂PO₄·H₂O (nominated as 0% dilution, serially diluted with ASW to 20, 35, 50 and 100% dilutions, 100% being pure ASW). The samples were pipetted into a plastic 1mL spectrophotometer cuvette with the clear detection window positioned perpendicularly to a camera lens. A white cardboard was placed behind the cuvette as background, and a picture was taken with a Canon EOS 7D MKII camera (Canon, Tokyo, Japan) equipped with a Laowa 100mm f/2.8 Ultra Macro APO 2:1 objective (Laowa, Hefei, China) and a speedlight flash. The pictures (jpeg, 5472 x 3648 pixels) were taken with 23 cm between the lens and the cuvette using the following settings: ISO 100, f/10, 1/100s exposure time. The flash intensity and direction were always the same.

The daily evaporation rate was estimated by comparing the ASW volumes inside the flasks at the end of the experimen and at the beginning of the experiment, taking into account the volume subtraction by the sampling process.

### Statistical analysis

The data were inspected using Shapiro-Wilk and Levene’s tests, and if the assumptions of normality and homogeneity of variances of parametric tests were met, the data were compared using two-tailed Student’s t-test (pairwise comparisons) or one-way ANOVA followed by *post hoc* Tukey’s test (multiple comparisons). When the assumptions of parametric tests were not met, either two-tailed Wilcoxon rank sum test (pairwise comparisons) or one-way Kruskal-Wallis followed by *post hoc* Dunn’s test (multiple comparisons) were used. The base significance level for all tests was p<0.05 and Bonferroni-corrected p-values were used in multiple comparisons. To assess whether the presence and absence of nutrients (phosphate or nitrogen) had an effect on the ΔF/F_M_’ of the sea slugs over the course of the starvation periods, the ΔF/F_M_’ values of the sea slugs were compared to their respective controls using a two-way ANOVA of aligned rank transformed data (two-way ART ANOVA). Two-way ART ANOVA was chosen, because individual sea slugs were not tracked through the time course of the starvation process, although repeated measurements were done in all sea slugs, preventing the use of traditional repeated-measures tests, like Repeated-Measures ANOVA or Linear Mixed-Effects Models. All statistical tests were done in R (v.4.2.3) using RStudio (2023.06.0+421; R Core Team, 2021). Figures were prepared with ggplot2 v3.4.1 package of R (Wickham, 2016) in conjunction with Inkscape (v.1.3). Non-outlier data in the box plots were calculated as ≤1.5 x the interquartile range.

### Ethics statement

This study was performed in accordance with EU legislation and directives concerning scientific research on animals, including the 3 R principles. Ethical approval is not required for studies conducted with non-cephalopod invertebrates. Only laboratory-reared animals were used in this work.

### Data availability

The source data will be made available upon publication after peer review.

## Results

### Kleptoplast longevity in *Elysia timida* increases in the presence of phosphate, but not in the presence of external nitrate or ammonium

A phosphate detection kit was used to get an estimate on whether *E. timida* sea slugs consumed phosphate from their surrounding media (containing 36 μM NaH₂PO₄·H₂O), and if the consumption was faster in the light than in the dark. All treatment flasks were aerated continuously, which did lead to evaporation rates of 1.33 ml/day ±SD 0.15 (control without sea slugs), 1.13 ±0.25 (sea slugs in the light) and 1.23 ±0.38 (sea slugs in the dark) even when the flasks were sealed with parafilm, only allowing the silicon air tube to pass through. Accordingly, phosphate concentration increased in the control flasks, based on the colorimetric analysis (Fig. 1). Phosphate concentration in the flasks containing *E. timida*, on the other hand, decreased both in the light and in the dark during four and seven of starvation, when compared to phosphate concentration in the control flasks of the same day, indicating that the sea slugs consumed phosphate from the surrounding media (Fig. 1).

**Figure 1.**
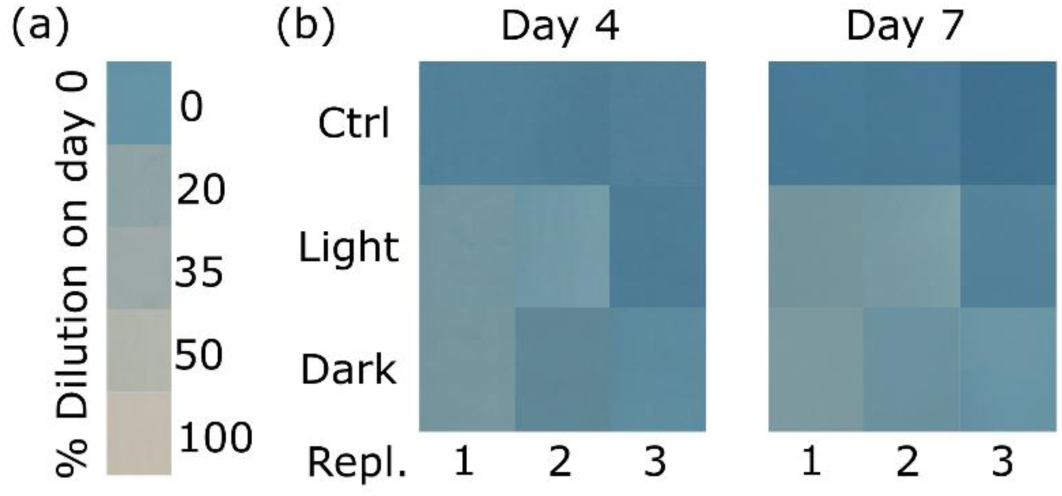
Phosphate uptake by *Elysia timida* in the light and in darkness. (a) Colorimetric detection of phosphates from artificial sea water (ASW) supplemented with 36 μM NaH₂PO₄·H₂O (0% dilution) and serially diluted with pure ASW. (b) Phosphates in control flasks without any sea slugs, in flasks kept in the light and containing 20 sea slugs per flask (Light) and in flasks kept in the dark and containing 20 sea slugs per flask (Dark) on days four and seven of starvation. The replicates have been measured from three separate flasks. The darkening blue color of the control flasks is likely the result of an increasing phosphate concentration caused by water evaporation from the flasks.

When *E. timida* individuals were starved in the presence of 36 μM NaH₂PO₄·H₂O (+P group), the PSII activity of their kleptoplasts (ΔF/F_M_’) remained higher through the 12-week starvation period than in the control group starved in ASW without any additions (Fig. 2a; individual effects of the treatment and the time in starvation were statistically significant both at p<0.001, as well as the combined effect of treatment x time in starvation at p<0.01, according to two-way ART ANOVA). The differences in ΔF/F_M_’ between the groups started to show at approximately weeks three or four, whereafter the PSII activity of the +P group remained higher until week 11; on week 12 there was very little PSII activity noticeable in either group. Sea slugs in the +P group did start to perish sooner (first deaths recorded on week seven) than in the control group (week eight), but after week eight the number of dead individuals was rising faster in the control than in the +P group (Fig. 2a). The final numbers of dead individuals after 12 weeks in starvation were 16 and 10 in the control and +P treatment groups, respectively. Rapid light curves (RLCs) were measured from the sea slugs on weeks zero and five (Fig. 2b,c). The maximum relative electron transfer rate of PSII (rETR_MAX_) was initially slightly lower in the +P group (Fig. 2b,d), but the differences were not statistically significant between the control and the +P group on week zero (two-tailed Student’s t-test, p=0.52). On week five, the rETR_MAX_ of the +P group was significantly higher than in the control group (Fig. 2c,d; two-tailed Student’s t-test, p<0.05).

**Figure 2.**
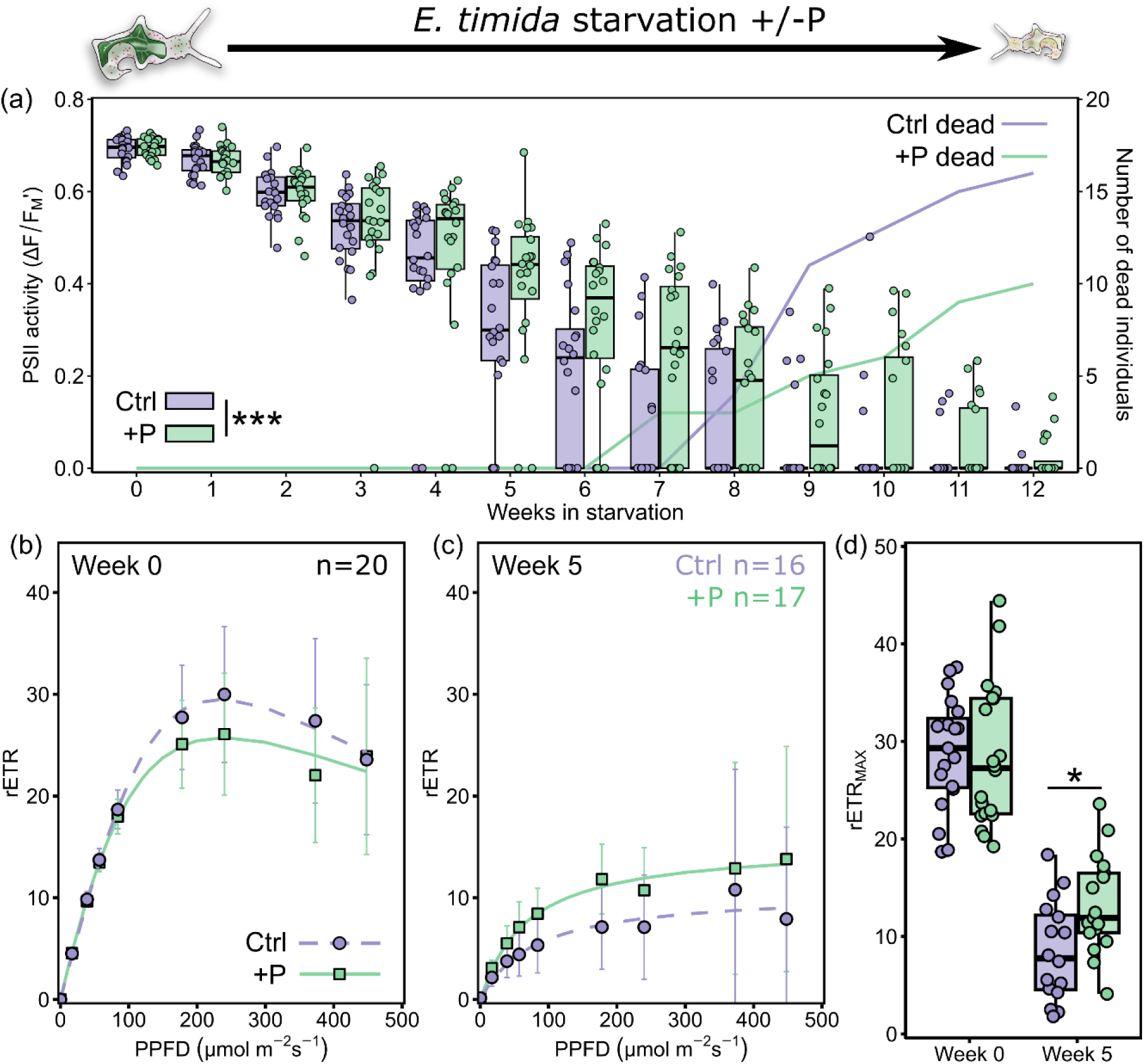
Photosynthetic activity of *Elysia timida* during starvation in the absence or presence of phosphate. (a) The decrease in PSII activity (ΔF/F_M_’; box plots) and the number of dead individuals (lines) during a 12-week starvation period in the absence (Ctrl; purple) and presence (+P; green) of 36 μM NaH₂PO₄·H₂O. Dead individuals were counted as having ΔF/F_M_’ of 0 (n=20 for the entire experiment for both groups). A significant difference in PSII activity caused by the treatment is indicated by *** (two-way ART ANOVA, p<0.001). (b, c) Relative electron transfer of PSII (rETR) during rapid light curve measurements from both groups (b) at the beginning of the experiment on week 0 and (c) on week 5 of starvation, modelled according to Eilers and Peeters (1988) (dashed and solid lines for Ctrl and +P, respectively). Only individuals where a reliable rapid light curve could be measured were used. The light treatment at each light intensity (photosynthetic photon flux density; PPFD) lasted for 60 s before a saturating light pulse was fired to estimate rETR. The data are means from the number of biological replicates shown in the panels. Error bars show standard deviation. (d) The rapid light curve parameter rETR_MAX_ (maximum relative electron transfer of PSII) in both treatments at week 0 and week 5 of the experiment. rETR_MAX_ was derived from modelling the data in panels (b, c). Significant differences between treatments are indicated by * (two-tailed Student’s t-test, p<0.05). The box plots in panels (a, d) show the medians and interquartile ranges, the whiskers indicate non-outlier maxima and minima, and each circle is an individual sea slug.

When the starvation experiment was repeated by starving the sea slugs in ASW without any additions (control group) and in ASW supplemented with 882 μM NaNO_3_ (+NaNO_3_) as a nitrogen source, there were no clear differences between the ΔF/F_M_’ values of the control and + NaNO_3_ group during a 10-week starvation period (Fig. 3a; two-way ART ANOVA, the effect of treatment and the combined effect of treatment x time in starvation were both nonsignificant at p=0.23 and p=0.85, respectively; the starvation time alone had a significant effect at p<0.001). The sea slugs started dying two weeks earlier in the +NaNO_3_ group (first deaths on week seven) than in the control group (on week nine), but the number of dead individuals, three and five for +NaNO_3_ and control, respectively, was rather similar at the end of week 10 (Fig. 3a). The RLCs measured on week zero and week five of starvation (Fig. 3b,c) did not show any statistically significant differences between the treatment groups, as compared based on the rETR_MAX_ parameter (Fig. 3d; Wilcoxon rank sum tests, on week 0 p=0.18, and on week 5 p=0.16).

**Figure 3.**
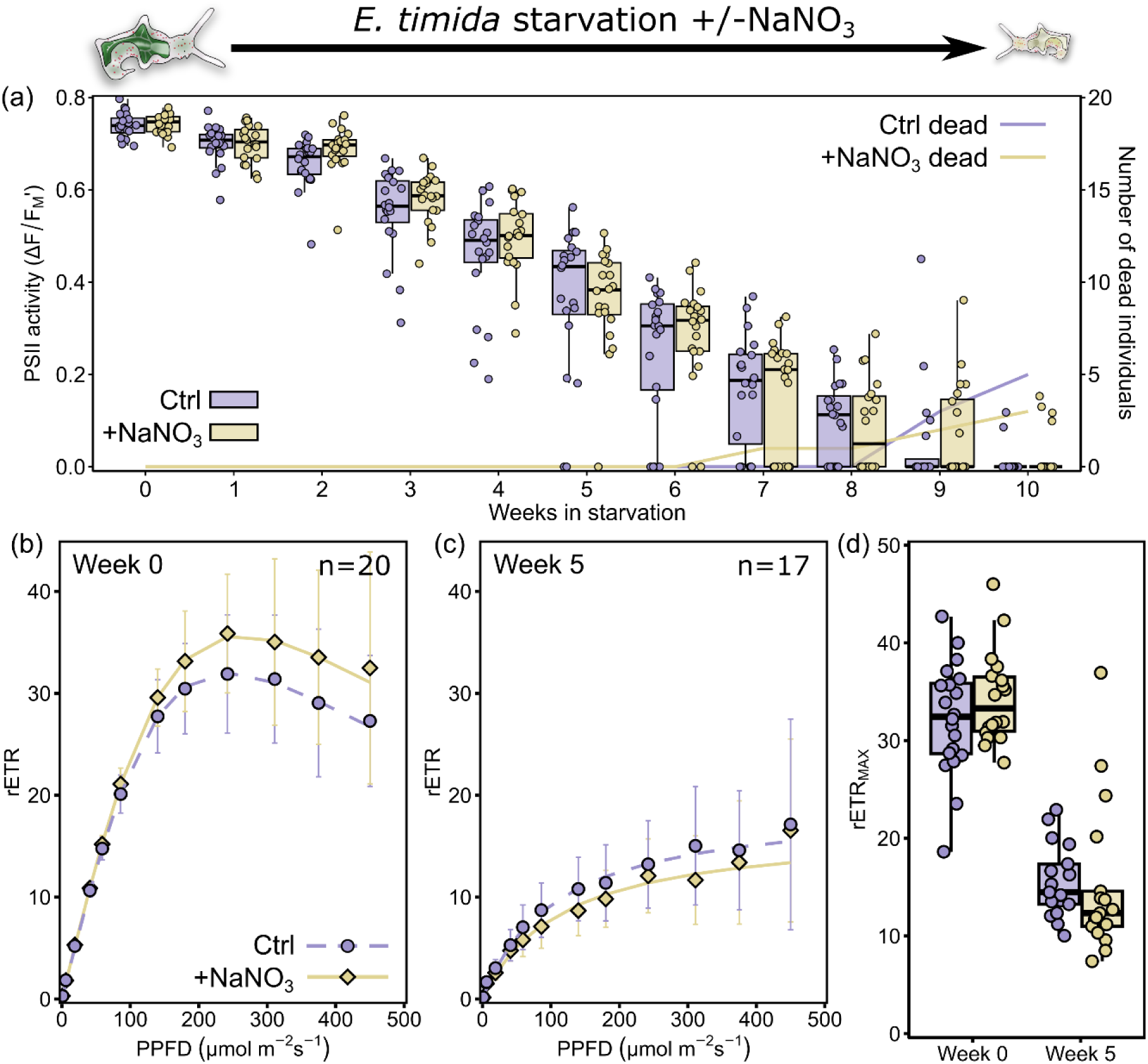
Photosynthetic activity of *Elysia timida* during starvation in the absence or presence of NaNO_3_. (a) The decrease in PSII activity (ΔF/F_M_’; box plots) and the number of dead individuals (lines) during a 10-week starvation period in the absence (Ctrl; purple) and presence (+NaNO_3_; yellow) of 882 μM NaNO_3_. Dead individuals were counted as having ΔF/F_M_’ of 0 (n=20 for the entire experiment for both groups). (b, c) Relative electron transfer of PSII (rETR) during rapid light curve measurements from both groups (b) at the beginning of the experiment on week 0 and (c) on week 5 of starvation, modelled according to Eilers and Peeters (1988) (dashed and solid lines for Ctrl and +NaNO_3_ respectively). Only individuals where a reliable rapid light curve could be measured were used. The light treatment at each light intensity (photosynthetic photon flux density; PPFD) lasted for 60 s before a saturating light pulse was fired to estimate rETR. The data are means from the number of biological replicates shown in the panels. Error bars show standard deviation. (d) The rapid light curve parameter rETR_MAX_ (maximum relative electron transfer of PSII) in both treatments at week 0 and week 5 of the experiment. rETR_MAX_ was derived from modelling the data in panels (b, c). The box plots in panels (a, d) show the medians and interquartile ranges, the whiskers indicate non-outlier maxima and minima, and each circle is an individual sea slug.

The effect of nitrogen on the longevity of the kleptoplasts was also tested by using 20 μM NH_4_Cl (+NH_4_Cl) as a nitrogen source (Fig. 4). NH_4_Cl had a clear negative effect on ΔF/F_M_’ of the kleptoplasts; ΔF/F_M_’ values started to decrease faster already in week one of the five-week starvation period in the +NH_4_Cl group compared to the control group, showing very little PSII activity by week five (Fig. 4a; statistically significant differences at p<0.001 by the treatment and by the time in starvation, as well as by the combined effect of treatment x time in starvation, according to two-way ART ANOVA). This was also reflected on the RLCs measured at week two of the starvation, as reliable RLCs could only be measured from 13 individuals up to PPFD 142 µmol m^-2^s^-1^ in the +NH_4_Cl group on week two, whereas all 20 individuals of the control group showed above zero rETR values up to PPFD 244 µmol m^-2^s^-1^ on the same week (Fig. 4b,c). On week zero, there were no statistically significant differences in the rETR_MAX_ parameter between the treatment groups (two-tailed Wilcoxon rank sum test, p=0.10), but on week two the rETR_MAX_ of the +NH_4_Cl group was significantly lower than in the control (Fig. 4d; two-tailed Wilcoxon rank sum test, p<0.05). There were no deaths in either of the groups during the five-week starvation period.

**Figure 4.**
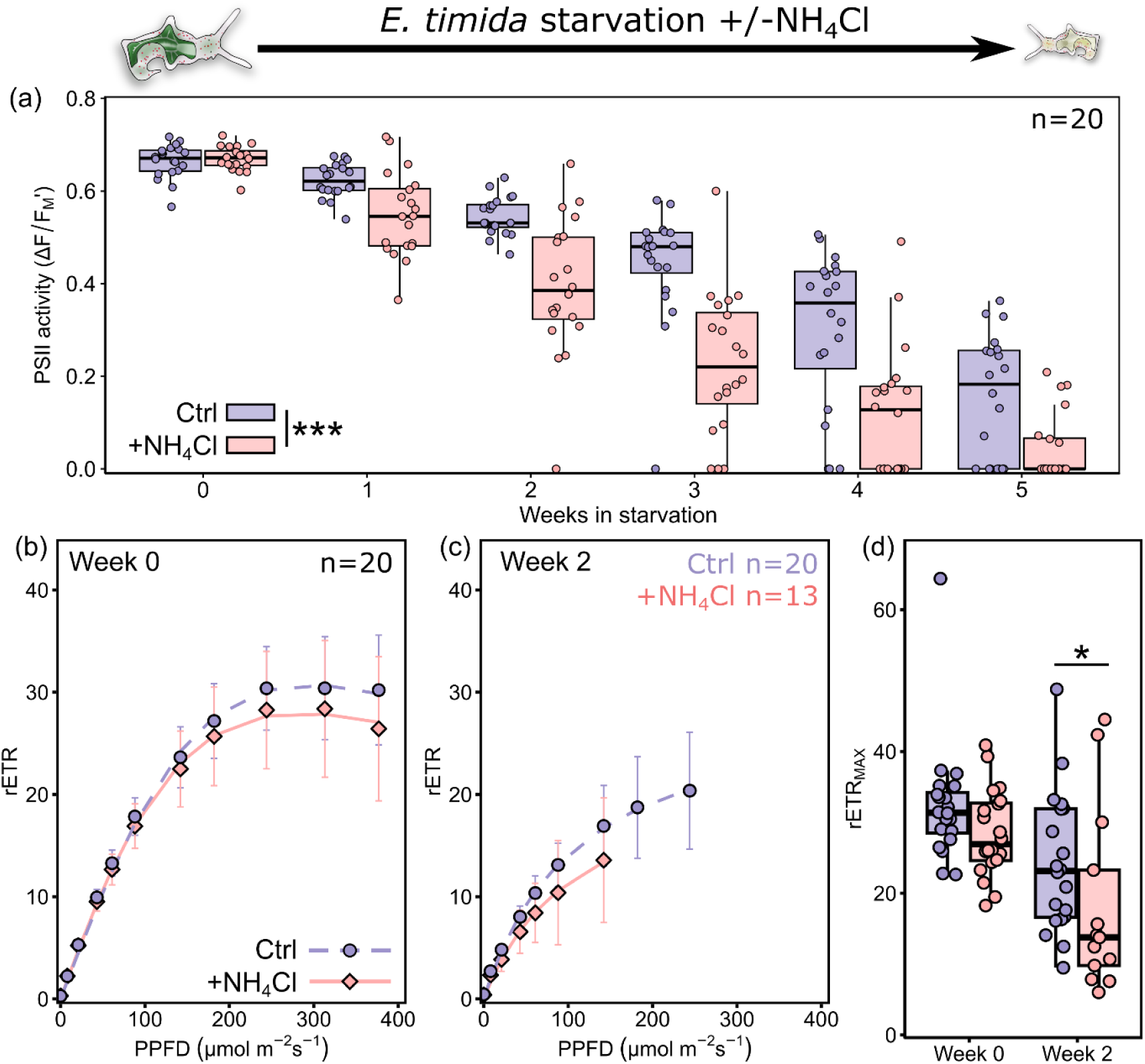
Photosynthetic activity of *Elysia timida* during starvation in the absence or presence of NH_4_Cl. (a) The decrease in PSII activity (ΔF/F_M_’; box plots) during a 5-week starvation period in the absence (Ctrl; purple) and presence (+NH_4_Cl; pink) of 20 μM NH_4_Cl. A significant difference in PSII activity caused by the treatment is indicated by *** (two-way ART ANOVA, p<0.001). (b, c) Relative electron transfer of PSII (rETR) during rapid light curve measurements from both groups (b) at the beginning of the experiment on week 0 and (c) on week 2 of starvation, modelled according to Eilers and Peeters (1988) (dashed and solid lines for Ctrl and +NH_4_Cl respectively). Only individuals where a reliable rapid light curve could be measured were used. The light treatment at each light intensity (photosynthetic photon flux density; PPFD) lasted 60 s before a saturating light pulse was fired to estimate rETR. The data are means from the number of biological replicates shown in the panels. Error bars show standard deviation. (d) The rapid light curve parameter rETR_MAX_ (maximum relative electron transfer of PSII) in both treatments at week 0 and week 2 of the starvation. rETR_MAX_ was derived from modelling the data in panels (b, c). Significant differences between treatments are indicated by * (two-tailed Wilcoxon rank sum test, p<0.05). The box plots in panels (a, d) show the medians and interquartile ranges, the whiskers indicate non-outlier maxima and minima, and each circle is an individual sea slug.

NH_4_Cl is a known uncoupler that disrupts the proton gradient across the thylakoid membrane. Because of this, we tested the short-term effect NH_4_Cl has on rETR and the induction of photoprotective non-photochemical quenching (NPQ) of excitation energy during a RLC in *E. timida* after 1 h incubation in 20 μM NH_4_Cl. The rETR and NPQ values were, however, nearly identical in the presence of NH_4_Cl compared to the control (only ASW) (Supplementary Fig. S2).

### Phosphate and nitrogen depletion in *Acetabularia acetabulum* leads to defective photosynthesis that is recoverable by nutrient repletion

The effect of phosphate and nitrogen deprivation to the photosynthetic electron transfer of the alga *A. acetabulum* was tested by growing them in their growth medium without NaH₂PO₄·H₂O (-P group) or NaNO_3_ (-N group). A control group of algae, grown in regular growth medium with all the nutrients, was monitored simultaneously through the 12-week nutrient depletion period (Fig. 5a-c). On week eight, the RLCs measured from the algae suggested that photosynthesis in the -N group was defective, as sensible non-zero rETR values could only be measured up to PPFD 85 µmol m^-2^s^-1^ (Fig. 5b). The rETR_MAX_ was also significantly lower in the -N group compared to the control group (one-way ANOVA, followed by *post hoc* Tukey’s test, p<0.01), whereas the rETR_MAX_ of the -P group was not statistically significantly different (p=0.30) from the control group, although already slightly lower. The nutrient depletion in the -P group took 12 weeks to clearly affect the photosynthetic electron transfer in *A. acetabulum*, when non-zero rETR could only be measured up to PPFD 85 µmol m^-2^s^-1^ and the rETR_MAX_ was statistically significantly lower in the -P group compared to the control (Fig. 5c; two-tailed Student’s t-test, p<0.05). The total amount of Chl per algae fresh weight in *A. acetabulum* depleted of nitrogen for eight weeks was 0.31 µg Chl mg^-1^ (SD±0.06, n=4), which was significantly lower than 0.51 µg Chl mg^-1^ (SD±0.06, n=4) measured from algae taken from growth conditions (with all f/2 nutrients) (two-tailed Student’s t-test, p<0.01). The -P group algae were not inspected for Chl contents due to limited amount of algae available for *E. timida* feeding experiments.

**Figure 5.**
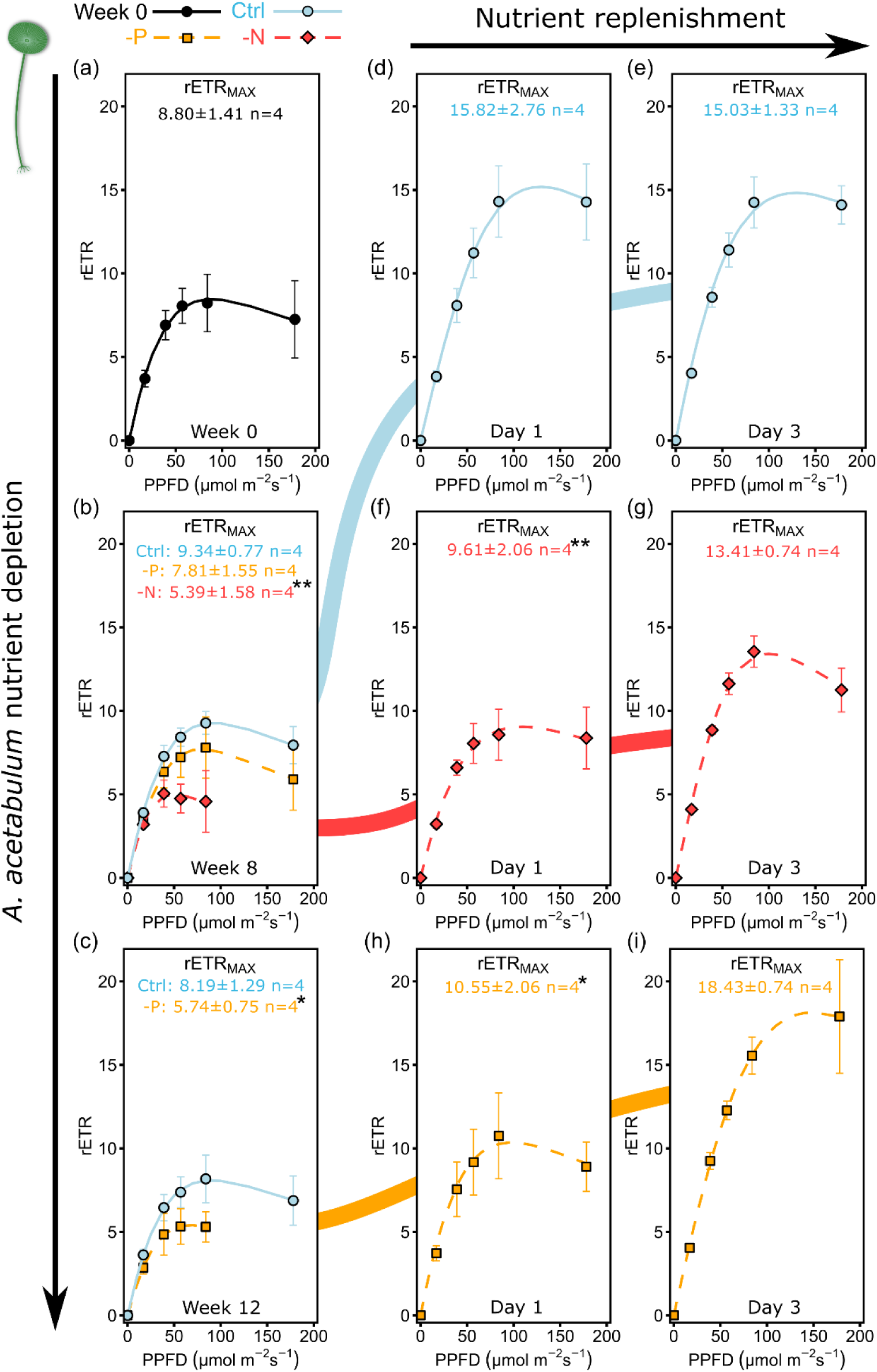
The photosynthetic effects and reversibility of phosphate and nitrogen depletion in the alga *Acetabularia acetabulum*. (a) Rapid light curve (RLC), measuring relative electron transfer of PSII (rETR) at different light intensities (photosynthetic photon flux density; PPFD), from algae at the start of the experiment on week 0. The light treatment at each light intensity lasted 60 s before a saturating light pulse was fired to estimate rETR. (b, c) RLCs from algae kept in full growth medium (Ctrl; blue) or in nutrient depleted growth media without either NaNO_3_ (-N; red) or NaH₂PO₄·H₂O (-P; orange) for (b) eight and (c) 12 weeks. The dashed and solid lines show the data modelled according to Eilers and Peeters (1988). Maximum rETR (rETR_MAX_) values derived from the modelled RLCs are shown in each panel (±standard deviation), and significant differences between Ctrl treatment of a specific week and either of the nutrient treatments of the same week are indicated by asterisks (one-way ANOVA and Tukey’s test in panel b, two-tailed Student’s t-test in c). (d-i) RLCs and rETR_MAX_ from (d, e) Ctrl, and (f, g) nitrogen and (h, i) phosphorus depleted algae after transfer to vials containing fresh growth media with all the nutrients, measured after one and three days of the replenishment. The coloured nodes in the background connecting the nutrient depletion panels (a-c) to the nutrient replenishment panels (d-i) show the week of nutrient depletion where the samples were taken for nutrient replenishment. Asterisks next to the rETR_MAX_ values in panels (d-i) indicate statistically significant differences between Ctrl treatment and either of the nutrient treatments of the same day (one-way ANOVA and Tukey’s test; * and ** indicate a significance at p<0.05 and p<0.01). All data are means from the number of biological replicates indicated in the panels. The error bars show standard deviation.

When part of the control group algae from week eight of the nutrient depletion experiment were placed in a separate smaller vial and given fresh growth medium (including all the nutrients), rETR increased immediately after one day in the replenishment conditions (Fig. 5d) and stayed at the same level until the end of the experiment on day three (Fig. 5e). The increase in rETR of the control group suggests that photosynthesis was limited even in the control group, even though a small portion of new growth media was added every week, perhaps due to the high density of algae. The rETR of the -N group algae taken from week eight of nutrient depletion also showed an initial increase after one day of nutrient replenishment with growth media containing all of the nutrients, although the rETR_MAX_ of the -N group was still significantly lower than in the control group on the same day of the replenishment (Fig. 5f; one-way ANOVA, followed by *post hoc* Tukey’s test, p<0.01 for Ctrl vs. -N). The rETR of the -N group further increased to similar levels as in the control group on day three of the replenishment (Fig. 5g). Finally, when the -P group algae that had been depleted of phosphate for 12 weeks were replenished with growth media containing all of the nutrients, they behaved similarly to the -N group algae, with the rETR_MAX_ increasing after one day, but still being statistically significantly lower than in the control group (Fig. 5h; one-way ANOVA, Tukey’s test, p<0.05 for Ctrl vs. -P). After three days in nutrient replenishment, the rETR_MAX_ was significantly higher in the -P group than in -N group (one-way ANOVA, *post hoc* Tukey’s test, p<0.05), but there were no statistically significant differences in either treatment group any more compared to the control group (Fig. 5i; Tukey’s test Ctrl vs. -N p=0.47, Ctrl vs. -P p=0.07).

The effects of nutrient depletion and replenishment on photosynthesis were also inspected in nitrogen depleted algae that were replenished with 20 µM NH_4_Cl, instead of NaNO_3_. The results show that *A. acetabulum* is also capable of using ammonium instead of nitrate as a source for nitrogen to recover from nitrogen depletion, as rETR completely recovered after the addition of NH_4_Cl (Supplementary Fig. S3). Finally, the Chl content of algae after three days of the replenishment with both NaNO_3_ (0.49 µg Chl mg^-1^ fresh weight, SD±0.03, n=4) and NH_4_Cl (0.44 µg Chl mg^-1^, SD±0.01, n=4), showed statistically significant increases compared to the nitrogen depleted algae (0.31 µg Chl mg^-1^, SD±0.06, n=4; one-way ANOVA and Tukey’s test, p<0.001 for depleted vs. replenished with NaNO_3_ and p<0.01 for depleted vs. replenished with NH_4_Cl).

### *Elysia timida* restores photosynthetic activity in phosphate-depleted kleptoplasts, but not in nitrogen-depleted ones

Phosphate and nitrogen (NaNO_3_) depleted algae from the nutrient depletion experiment (shown in Fig. 5b,c) were used to test if incorporation of chloroplasts as kleptoplasts by the sea slug *E. timida* restores their photosynthetic activity back to non-depleted levels. For this, *E. timida* individuals that had been bleached of their kleptoplasts using a high light treatment were re-fed by supplying them specifically with control algae, -N group algae after week eight of depletion (Fig. 5b) or -P group algae after week 12 of depletion (Fig. 5c). The sea slugs were allowed to continuously feed on the algae in a large 150 L life support system (LSS) filled with ASW. RLCs were measured from the control, -N and -P groups (named after the algae they were feeding on) after one, three and seven days of feeding (Fig. 6a-c), and the rETR_MAX_ parameters derived from the RLCs were used for statistical comparison of the groups (Fig. 6d-f).

**Figure 6.**
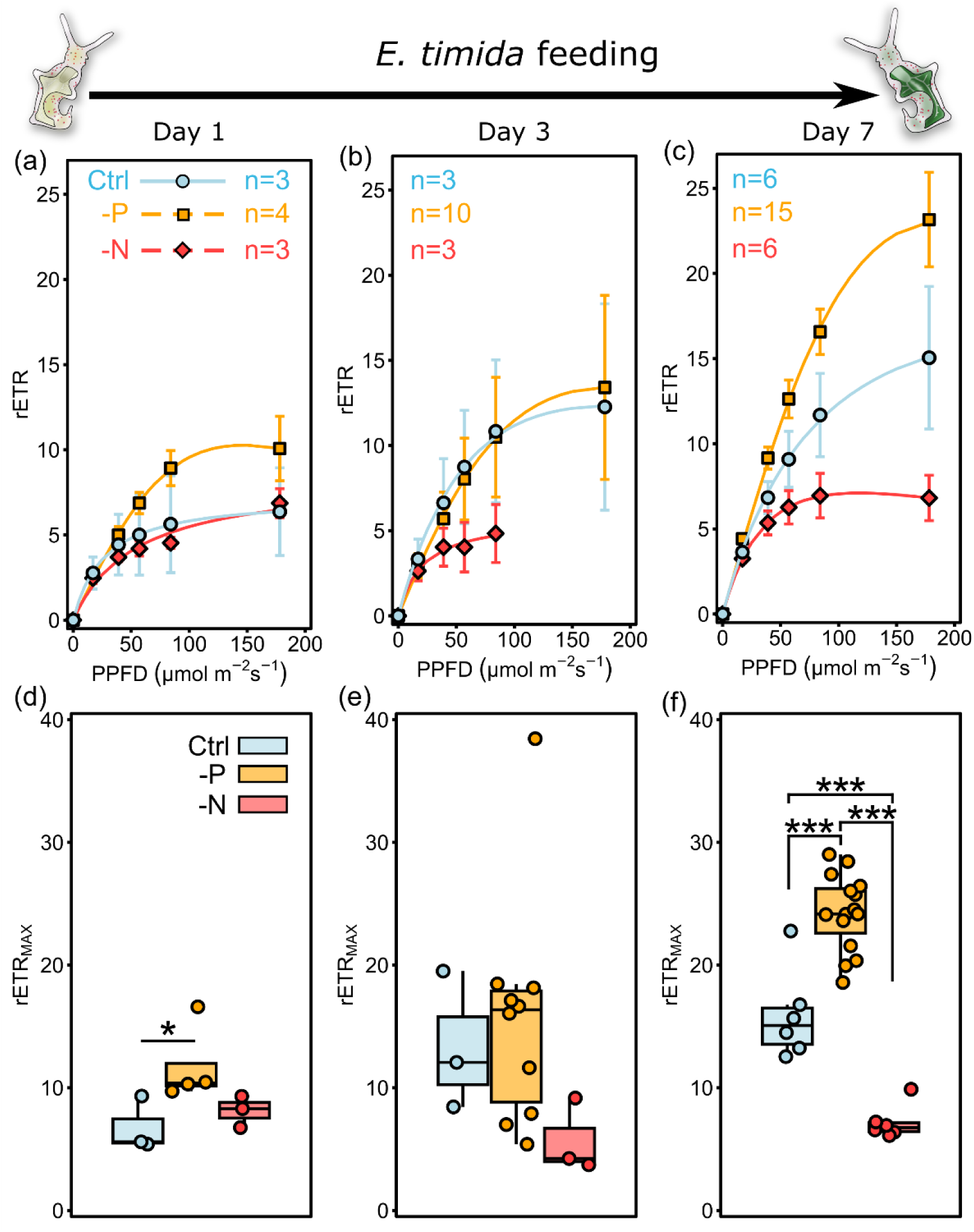
Photosynthetic recovery of nutrient deprived kleptoplasts in the sea slug *Elysia timida* in the absence of external nutrients. (a-c) Relative electron transfer of PSII (rETR) during rapid light curves (RLCs) at different light intensities (photosynthetic photon flux density; PPFD) measured from bleached sea slugs after (a) one, (b) three and (c) seven days of feeding on control (Ctrl; blue) and nitrogen (-N; red) or phosphate deprived (-P; orange) algae. For the rETR values of the algae (prior to feeding to the sea slugs), see Fig. 5a-c. All data are means from the number of biological replicates indicated in the panels. The error bars show standard deviation, and the lines show the data modelled according to Eilers and Peeters (1988). (d-f) Maximum rETR (rETR_MAX_), derived from the modelled curves in (a-c), in the different feeding groups after (d) one, (e) three and (f) seven days of feeding. The box plots show the medians and interquartile ranges, whereas the whiskers indicate non-outlier maxima and minima. Each circle in the boxplots is an individual sea slug. Significant differences between the treatments are indicated by asterisks (Kruskal-Wallis and Dunn’s test in panel d,e, one-way ANOVA and Tukey’s test in f; * and *** indicate significant differences at p<0.05 and p<0.001, respectively).

After one day of feeding, the control and -N groups of sea slugs exhibited very similar RLCs to each other, whereas the -P group showed fastest rETR out of all groups (Fig. 6a), and the rETR_MAX_ of the -P group was also statistically significantly higher than in the control group (Fig. 6d; Kruskal-Wallis and Dunn’s test, p<0.05). After three days of feeding, the RLCs of the control and the -P groups were similar, both showing an increasing trend in rETR compared to day one of feeding (Fig. 6b). The photosynthesis of the -N sea slugs used in the measurements on day three of feeding was clearly defective, as the RLCs showed non-zero rETR values only up to PPFD 85 µmol m^-2^s^-1^, unlike in the control and -P groups (Fig. 6b), but there were no statistically significant differences between the rETR_MAX_ of the groups (Fig. 6e; Kruskal-Wallis, p=0.11). After seven days of feeding the differences between the treatment groups became very clear (Fig. 6c). The rETR of the -P group sea slugs exceeded the rETR of the alga *A. acetabulum* measured in any of the experiments of this study (Fig. 5), and the rETR_MAX_ of -P sea slugs was also statistically significantly higher than in either of the other two treatment groups on day seven (Fig. 6f; one-way ANOVA and Tukey’s test, p<0.001), whereas the RLC of the control group sea slugs (Fig. 6c) resembled the RLCs measured from the algae after nutrient replenishment (Fig. 5d,e). In the -N group of sea slugs, the rETR remained low even on day seven of feeding, although photosynthesis could be estimated up to PPFD 180 µmol m^-2^s^-1^ (Fig. 6c), unlike on day three. The rETR_MAX_ of the -N group was also significantly lower than in the control group on day seven (Fig. 6f; one-way ANOVA and Tukey’s test, p<0.001).

### Photosynthetic activity in phosphate-depleted kleptoplasts decreases rapidly in the absence of phosphate uptake by the sea slugs

Next, the sea slugs that had been feeding on phosphate-depleted algae for seven days (Fig. 6c,f) were taken away from their food source and starved for 11 days in ASW without any added phosphate (-P starvation group) or in the presence of 36 μM NaH₂PO₄·H₂O (+P starvation group) (Fig. 7). The RLCs showed a more rapid decline in rETR in the -P starvation group compared to the +P starvation group (Fig. 7a,b). PSII activity during starvation (estimated with ΔF/F_M_’) differed statistically significantly between the -P and +P starvation groups only after one day in starvation (two-tailed Student’s t-test, p<0.05), even though ΔF/F_M_’ values were slightly higher in the +P group on both days one and 11 (Fig. 7c). However, the rETR_MAX_ parameter was statistically significantly higher in the +P group after 11 days in starvation (two-tailed Student’s t-test, p<0.05), while on day one rETR_MAX_ did not show significant differences (Fig. 7d; Wilcoxon rank sum test p=0.46). One sea slug from the -P group died during the 11-day starvation period, whereas there were no deaths in the +P group. Similar, but not as strong, results were observed when the experiment was repeated with more replicates and a longer time frame of the experiment, and the bleached sea slugs were fed with seven-weeks phosphate-depleted algae (instead of 12-weeks) and starved for five weeks (Supplementary Fig. S4).

**Figure 7.**
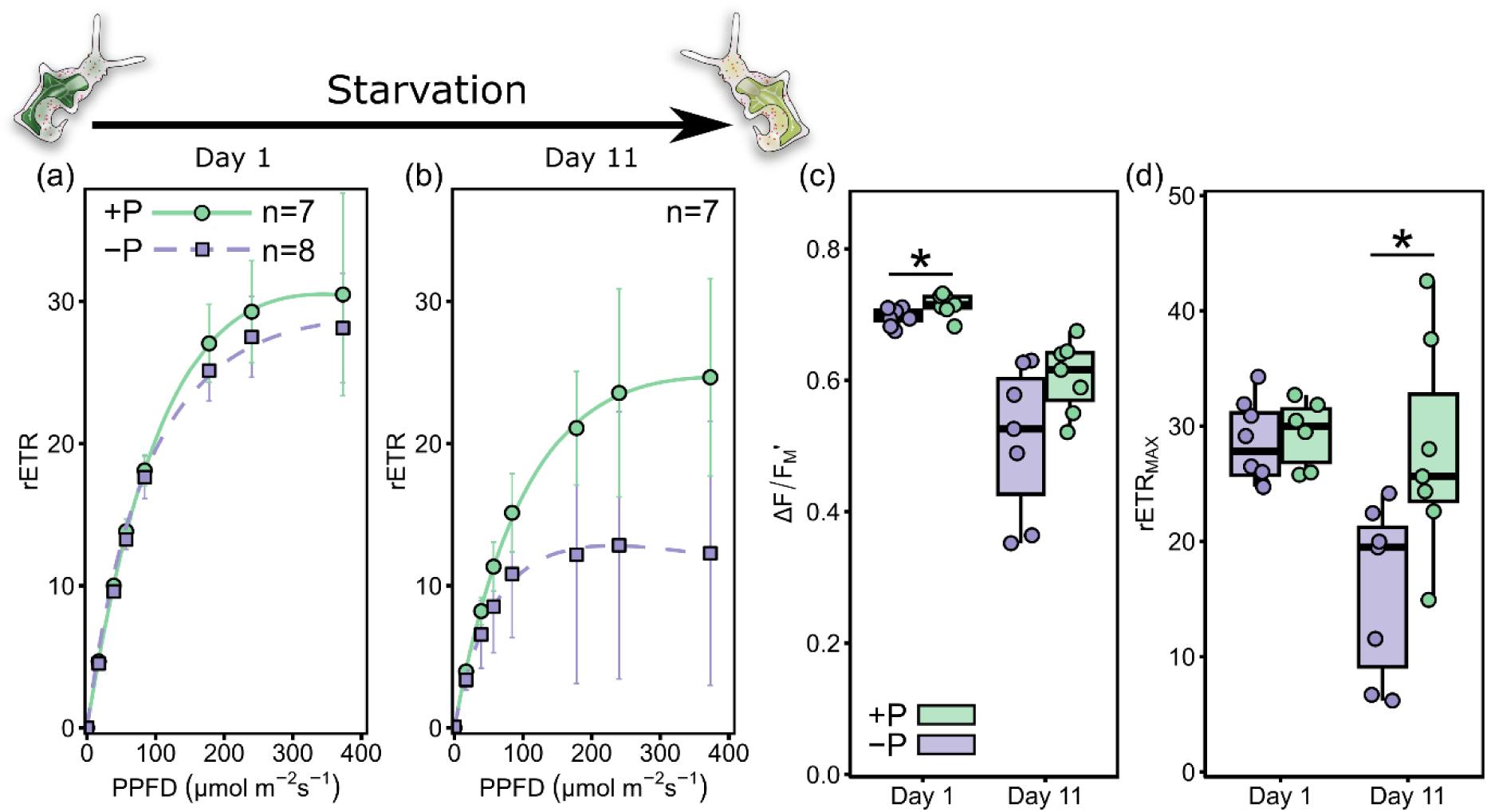
Short-term effect of added phosphorus on the photosynthesis of phosphorus deficient kleptoplasts in the sea slug *Elysia timida*. Rapid light curves (RLC) at different light intensities (photosynthetic photon flux density; PPFD) measured from sea slugs fed with phosphorus deprived algae (see Fig. 5c) and put to starve in the presence (+P; green) and absence (-P; purple) of phosphate (36 μM NaH₂PO₄·H₂O) for (a) one and (b) 11 days. (c) ΔF/F_M_’ and the rapid light curve parameter (d) rETR_MAX_ (maximum relative electron transfer of PSII) derived from the RLC data in panels (a,b) after modelling according to Eilers and Peeters (1988) (* and ** indicate significant differences at p<0.05; two-tailed Student’s t-test or two-tailed Wilcoxon rank sum test, respectively). The data points in (a,b) are means from the number of biological replicates used in the RLC measurements shown in panels. The lines show the modelled data and error bars in panels (a,b) indicate standard deviations. The box plots in (c,d) show the medians and interquartile ranges, whereas the whiskers indicate non-outlier maxima and minima. Each circle in the boxplots is an individual sea slug.

The rETR_MAX_ from the first week (measured on day 13 of starvation; note that the time course starts from week 0) of the first starvation experiment where the sea slugs were fed with non-depleted algae and starved in the presence and absence of phosphates (Fig. 2) was 30.18 (SD± 4.25, n=20) and 30.45 (± 6.31, n=20) for the -P and +P groups, respectively, whereas the rETR_MAX_ of the -P and +P sea slugs from individuals starved for 11 days (Fig. 7) was 15.79 (± 7.51, n=7) and 27.95 (± 9.33, n=7). Statistical comparison of rETR_MAX_ values from these two different starvation experiments showed significant differences between the -P groups (Kruskal-Wallis and Dunn’s test, p<0.001), whereas there were no differences between the +P groups (Dunn’s test, p>0.99). Thus, whether the sea slugs were fed with phosphate depleted or non-depleted algae, the rETR stayed high if external phosphate was provided, but a severe 12-week phosphate depletion of the algae prior to kleptoplast incorporation and starvation made the kleptoplasts very susceptible to degradation in the absence of external phosphate.

## Discussion

### *Elysia timida* enhances the longevity of kleptoplasts by providing them with phosphates

A study by Cobb (1978) showed that the kleptoplastic sea slug *Elysia viridis* accumulates polyphosphates in its tissues from inorganic phosphates, and that the accumulation of polyphosphates in this animal is likely due to their assimilation specifically in the kleptoplasts. We confirmed the previous results on phosphate intake by kleptoplastic sea slugs (Cobb, 1978) showing that the sea slug *E. timida* takes in phosphate both in the light and in the dark (Fig. 1). Despite showing that phosphates are taken in from the surrounding media by the sea slugs and that they seem to get transported to the kleptoplasts, Cobb (1978) did not discuss their importance for maintaining photosynthesis, rather, the author focused on the possibility that the sea slugs use the kleptoplasts as phosphate assimilators that provide the sea slugs with additional phosphates in times of starvation. Our results show that the addition of phosphates to the surrounding media of *E. timida* individuals during starvation has a strong positive effect on the longevity of their kleptoplasts, noticeable in the effective PSII activity parameter ΔF/F_M_’ (Fig. 2a) as well as in the relative electron transfer rate (rETR_MAX_) of PSII (Fig. 2b-d).

Like all animals, the sea slugs also use phosphates in their own non-kleptoplast related cell chemistry, and the added phosphates could be affecting the longevity of the kleptoplasts indirectly; a sea slug with abundant nutrients could be less inclined to degrade the kleptoplasts for their nutritive contents. The number of dead individuals was indeed lower in the +P starvation group (10) than in the control (16) at the end of the 12-week starvation period, which suggests that the +P group individuals were in generally better condition. On the other hand, the +P group sea slugs started to die one week earlier than the control animals (Fig. 2a), making it difficult to assess the general health benefits of added phosphates.

Evidence supporting a direct role of phosphates in keeping photosynthesis functional are provided by the results from the feeding experiment, where we first depleted *A. acetabulum* of phosphates until clear defects in the photosynthetic electron transfer were observed (Fig. 5c), after which the phosphate depleted algae were fed to *E. timida* individuals that were previously bleached of their old kleptoplasts (Fig. 6). *E. timida* showed a remarkable capacity to recover the low rETR_MAX_ phenotype of phosphate depleted kleptoplasts even in the absence of additional phosphates in the surrounding medium (Fig. 6), and the recovery was similar to or even stronger than in the phosphate depleted algae themselves when they were replenished with phosphate containing media (Fig. 5c,h,i). The results support the view that the sea slugs are indeed providing the kleptoplasts with extra phosphate from their own cellular reserves. The result that similar recovery was not observed in nitrogen depleted algae (Fig. 6) suggests that *E. timida* is particularly capable of aiding phosphate depleted kleptoplasts and that phosphate directly helps maintain photosynthetic electron transfer. The phosphate intake possibly allows photophosphorylation to produce ATP for carbon fixation, which in turn produces a strong electron sink in the kleptoplasts, decreasing the electron pressure and concomitant oxidative stress in the photosynthetic electron transfer chain (Dietz & Foyer, 1986).

Recently, *E. timida* and other kleptoplastic sea slugs were shown to induce strong structural changes to the kleptoplasts during their incorporation (Havurinne et al. 2025). These changes coincide with alterations to the light harvesting complexes of the kleptoplasts, that can also lead to increased photosynthetic electron transfer in some cases (Havurinne et al. 2025). Therefore, we considered the possibility that the structural changes of the kleptoplasts during incorporation are behind the recovery of rETR_MAX_ (Fig. 6), but if they were, they would have to enhance the photosynthesis of the control, phosphate and nitrogen depleted kleptoplasts of the sea slugs differently to explain the differences in the rETR_MAX_ of the sea slugs from the different treatment groups of the feeding experiment. Therefore, a synergistic effect of structural changes and nutrient delivery to the kleptoplasts is a more plausible explanation. The most notable change observed by Havurinne et al. (2025) was the immediate enforcement of light harvesting complex of PSII (LHCII) to serve PSII instead of PSI during kleptoplast incorporation. Interestingly, the movement of LHCII between PSII and PSI, i.e. state transitions, is controlled by phosphorylation of LHCII (Longoni et al. 2015). Our present results show that the sea slugs do take phosphate in, suggesting that the observed lack of state transitions in the kleptoplasts is not due to phosphate deficiency, but rather due to preferential use of phosphate to boost photosynthesis by other means in the kleptoplasts.

When the sea slugs that had been fed with phosphate depleted algae for seven days were placed to starve for 11 days in the presence and absence of phosphate, the decrease in rETR_MAX_ in the -P group was very fast compared to the +P group (Fig. 7), and also compared to the -P group of the first starvation experiment where the kleptoplasts derived from non-depleted algae (Fig. 2). This supports the earlier suggestions that the sea slugs mediate phosphate to the kleptoplasts from their surroundings (Fig. 2) and their own cellular reserves (Fig. 6), but it also suggests that when the kleptoplasts are already severely compromised in their phosphate requirements, the need for extra phosphate from the surrounding media to maintain photosynthesis exacerbates. It should be noted that extra phosphate in the media is certainly not a panacea that automatically helps maintain active photosynthesis forever, especially in sea slugs that have gone through a starvation and high light treatment that eliminates their old kleptoplasts. Indeed, when we replicated the experiment shown in Fig. 7 with more robust population sizes for both +/-P groups, this time feeding the bleached sea slugs with only seven-weeks phosphate-depleted algae (instead of 12-weeks like before in Fig. 7), the results once again showed that the ΔF/F_M_’ and rETR_MAX_ were positively affected by extra phosphate, but both groups showed very little photosynthetic activity by week five in starvation (Supplementary Fig. S4). Whether the starvation and high light bleaching affect the sea slugs’ capacity to maintain the kleptoplasts was not tested here.

It is still unclear which mechanisms are in place in photosynthetic sea slugs to mediate the uptake of exogenous phosphate and transport it to the kleptoplasts. In another non-kleptoplastic marine sea slug, *Aplysia californica*, a sodium/phosphate symporter seems to be involved in phosphate uptake (Gerencser et al. 2002), which has also been shown to be relevant for phosphate uptake in corals (Godinot et al. 2011). It is therefore plausible that similar symporters exist in sacoglossan sea slugs as well, and the recent proteomic inspection of the kleptosome, the organelle hosting the kleptoplasts in the sea slug *E. crispata*, provided evidence for a sodium-dependent phosphate transport protein 2B in the kleptosome (Allard et al. 2025). The phosphate transporters of green algae and their chloroplasts are not well known even in model species, not to mention in little studied green macroalgae like *A. acetabulum* or other members of the Ulvophyceae. With the current knowledge, it can only be speculated that like the green microalga *Chlamydomonas reinhardtii*, *A. acetabulum* chloroplasts may contain PHT4 family transporters in their envelope membranes that can facilitate phosphate intake to the chloroplasts (Tóth et al. 2024), and therefore also to kleptoplasts.

### Exogenous nitrate or ammonium are not beneficial for kleptoplast longevity

In the light, the kleptoplasts of the sea slugs *E. timida* and *E. viridis* are likely involved in assimilating nitrogen from ammonium, presumably to the amino acid glutamate via the glutamine synthetase (GS) and glutamate synthase (GOGAT) pathway present in algae chloroplasts (Teugels et al. 2008; Cruz et al. 2020; Cartaxana et al. 2021). The sea slugs do also assimilate some nitrogen in the dark, and previously this has been attributed to the mitochondrial glutamine dehydrogenase (GDH) pathway (Cruz et al. 2020; Cartaxana et al. 2021). However, a recent transcriptomic analysis of *E. viridis* showed that the sea slugs possess also their own GS and GOGAT enzymes that were upregulated when the animals were starved in the light and in the dark, suggesting that GDH is not the only nitrogen assimilation pathway of these animals (Frankenbach et al. 2023). This draws interesting parallels between the kleptoplastic sea slugs and symbiotic Aiptasia sea anemones; the host GS-GOGAT pathway has been suggested to be a major contributor to nitrogen cycling in Aiptasia in addition to the GS-GOGAT of their symbiotic algae (Cui et al. 2023). GOGAT is not universal in animals, but it has been shown to contribute to nitrogen cycling also in nematodes (Umair et al. 2011), and the enzyme seems to be present in certain insects as well (Hirayama et al. 1998). Importantly, the GOGAT of animals uses NAD(P)H as reducing power for nitrogen assimilation (Hirayama et al. 1998; Lehnert et al. 2014), whereas the chloroplastic GOGAT uses ferredoxin (Fd), connecting the Fd-GOGAT directly to photosynthetic electron transfer from PSI (Yang et al. 2016).

In *E. viridis,* assimilation of nitrogen has also been shown to take place when the animals are exposed to nitrite and urea (Teugels et al. 2008). These findings indicate that the sea slugs do take in nitrogen in multiple forms, but they also suggest that the hosts may be able to recycle their own nitrogenous waste products into useful biomolecules using their kleptoplasts to benefit themselves and the longevity of the kleptoplasts. Indeed, experiments using radiolabeled amino acids have shown that the kleptoplasts of the sea slug *E. chlorotica* do maintain *de novo* synthesis of chloroplast-encoded proteins even after months of starvation, including the D1 protein of PSII (Mujer et al. 1996). However, like with the phosphates, the effect of exogenous nitrogen sources on kleptoplast longevity was not previously evaluated.

In algae, nitrogen assimilation from nitrate requires the reduction of nitrate to nitrite by the cytosolic nitrate reductase. Nitrite is then transported to the chloroplast via transporters, like the NAR1 family of transporters in *C. reinhardtii* (Rexach et al. 2000), whereafter nitrite forms ammonium after reduction by the nitrite reductase in the chloroplast, and ammonium is assimilated in the GS-GOGAT pathway (Fernandez & Galvan, 2008). In line with this, Teugels et al. (2008) showed that *E. viridis* does not assimilate nitrogen from nitrate, because the animals do not have access to the cytosolic nitrate reductase of the algae. Sacoglossan sea slugs *Elysia rufescens* and *E. chlorotica* have been shown to contain a diverse bacterial gut microbiome, and many of the identified bacteria associated with the sea slugs have the capacity to reduce nitrate (Devine et al. 2012; Davis et al. 2013), possibly making it also accessible to the animals in long-term starvation experiments. The microbiome of our *E. timida* lab population is unknown, but our results show that additional nitrate did not influence the longevity of its kleptoplasts in starvation (Fig. 3), suggesting that even if the gut bacteria were able to assimilate trace amounts of nitrogen from nitrate, it is negligible in terms of kleptoplast functionality.

In water, ammonium and the unionized form ammonia, NH_3_, exist in equilibrium that is affected by salinity, temperature and pH (Bell et al. 2008). Ammonia is more harmful, but both can interfere with many cellular functions, and they are toxic in high concentrations, affecting, for example, the osmotic regulation of aquatic organisms (Lin et al. 1993). For example, the exposure of the shrimp *Litopenaeus vannamei* to elevated levels of either ammonium or ammonia can eventually lead to elevated oxidative stress and trigger mitochondrial dysfunction and apoptosis (Tong et al. 2024). The toxic effects of ammonium/ammonia on marine animals are usually tested by exposing them to different concentrations of these chemicals for hours or days and then analyzing the stress markers, including mortality, during the exposure (Lin et al. 1993; Tong et al. 2024). The concentrations of NH_4_Cl used in these tests vary a lot, but the concentration used in the present study, 20 µM, is a relatively high concentration of NH_4_Cl, especially for a five-week long exposure of the animals. It is therefore surprising that during the starvation experiment of the sea slug *E. timida* in the presence and absence of 20 µM NH_4_Cl (Fig. 4) there were no deaths in either group at the end of the experiment after five weeks, suggesting that *E. timida* has a high tolerance against the toxic effects of ammonium.

The photosynthetic activity of the kleptoplasts during starvation decreased faster in the presence of ammonium than in the control group. The faster decay of ΔF/F_M_’ was already noticeable after one week in starvation (Fig. 4a), and photosynthetic electron transfer was difficult to assess from the +NH_4_Cl group after two weeks due to the poor Chl fluorescence signal from many of the individuals, unlike in the control group (Fig. 4c,d). While the animals themselves might be resistant against ammonium toxicity, it is possible that the kleptoplasts are not. Ammonium/ammonia are also known to interfere with photosynthesis, resulting in damage to PSII, presumably by ammonium disrupting the functionality of the oxygen evolving complex of PSII through water ligand replacement (Tsuno et al. 2011; Wang et al. 2019). NH_4_Cl at millimolar concentrations is also commonly used as an uncoupler of photophosphorylation in algae and plants, as it disrupts the proton gradient across the thylakoid membrane and can lead to problems in ATP production. The uncoupling can also lead to defects in the induction of NPQ (Sherman & Wimmer, 1983; Blommaert et al. 2021). However, based on the very similar NPQ levels during RLCs between the control and after 1 h incubation in 20 µM NH_4_Cl (Supplementary Fig. S2), the negative effects of ammonium to kleptoplast longevity are not likely due to uncoupler effects. The kleptoplasts in *E. timida* are capable of assimilating NH_4_Cl (Cartaxana et al. 2021), but due to the problematic side effects of ammonium, we can only conclude that the addition of 20 µM NH_4_Cl is not beneficial for kleptoplast longevity. Importantly, this does not mean that nitrogen provided by the animal from its own reserves is not important to the kleptoplasts.

Eight-weeks nitrogen-depleted *A. acetabulum* algae were able to recover their photosynthetic activity in three days when replenished with f/2 medium containing either nitrate (Fig. 5b,f,g) or ammonium (Supplementary Fig. S3) as a nitrogen source, and the recovery was even faster with the relatively high NH_4_Cl concentration of 20 µM. This suggests that in the algae this concentration is below the toxic level, at least in terms of affecting photosynthesis, and the algae can reverse the effects caused by nitrogen depletion using multiple nitrogen sources. The toxic effects of ammonium to the photosynthesis in kleptoplasts of the sea slugs may be exacerbated by the fact that the animal cell environment is expected to contain far more nitrogenous waste products, including ammonium, than the algae even without administration of extra nitrogen. However, unlike sea slugs that were fed phosphate depleted algae, *E. timida* individuals fed with nitrogen depleted algae were not able to recover the photosynthetic capacity of their kleptoplasts during seven days of feeding (Fig. 6). The defects caused by nitrogen depletion were therefore more severe than those caused by phosphate depletion, and the repair processes would likely require input from nucleus encoded proteins of the algae.

Nitrogen is a major component of Chl, and the defining steps of Chl synthesis take place in chloroplasts, but the key enzymes of the synthesis pathway are mostly nucleus encoded in plants and algae (Bryant et al. 2020; Zhao et al. 2020). Two reports of Chl synthesis in kleptoplastic systems exist; one in *E. chlorotica*, where exogenous labeled 5-aminolevulinic acid (a precursor of Chl) was shown to result in radiolabeled Chl *a* in the animal (Pierce et al. 2009), and one in the kleptoplastic ciliate *Mesodinium rubrum* that was shown to be able to increase the amount of Chl *a* in its cells in low light conditions (Johnson et al. 2023). However, the genes proposed to have been horizontally transferred into *E. chlorotica* genome that could facilitate Chl synthesis in the animal have not been verified in later genomic or transcriptomic analyses (Bhattacharya et al. 2013; Chan et al. 2018; Cai et al. 2019; Maeda et al. 2021), and *M. rubrum* also temporarily sequesters the nucleus (kleptokaryon) from its prey, which does contain the genes for Chl synthesis (Johnson et al. 2023). Therefore, it seems evident that Chl synthesis in kleptoplasts is an exception, not a rule. A common response to nitrogen deficiency in algae is a decrease in the amount of Chl (Fattore et al. 2021), which was also observed here in *A. acetabulum* after eight weeks in nitrogen depletion. While the nitrogen depleted algae were able to resynthesize Chl after nitrogen repletion using either nitrate or ammonium, we suggest that the incapability of *E. timida* to synthesize Chl is one of the fundamental reasons why the sea slugs were not able to recover the photosynthetic activity of nitrogen depleted kleptoplasts. Even if the nitrogen provided by the host to the kleptoplasts would in theory allow them to synthesize some proteins of the photosynthetic machinery to repair it, many of those proteins, like the D1 protein of PSII, would still require Chl to assemble and function properly (Komenda et al. 2024).

### Synergistic effects of phosphate and nitrogen on kleptoplasts

In symbiotic systems like corals, the importance of phosphate is often discussed in relation to the amount of available nitrogen. A balanced supply ratio of phosphate and nitrogen, mandated by the organisms and their environment, is required for optimal algal growth and maintaining a healthy, mutualistic relationship between a coral host and its symbionts (Rosset et al. 2017; Morris et al. 2019; Yaakob et al. 2021). The symbiotic zooxanthellae of corals are living organisms that can grow and divide inside the host, therefore possessing multiple ways of storing nutrients provided by the host. This is in stark contrast to the kleptoplasts inside a sea slug; kleptoplasts do not divide and are expected to have less options for nutrient allocation compared to their original location as chloroplasts inside the algal cell.

Although no data exists on the topic, it is plausible to assume that due to the differences in nitrogen metabolism between algae and kleptoplastic sea slugs, the amount of nitrogenous waste products like ammonium in the cellular environment of the kleptoplasts increases to higher levels than what is usually present for chloroplasts in free-living algae. Most starvation experiments with the sea slugs have been done in artificial sea water that contains only trace amounts of phosphates and nitrogen sources (e.g., Green et al. 2000; Laetz & Wägele, 2018; Cartaxana et al. 2023), which may expose the kleptoplasts inside the animals to an environment containing plenty of nitrogen (e.g. ammonium; due to the waste metabolism of the slug) but relatively limited free phosphates, which the sea slugs themselves may need to maintain their own cellular functions. The sea slugs could still supply the kleptoplasts with also phosphate during times when nutrients are not scarce, like during feeding (Fig. 6) and at the beginning of starvation. When starvation proceeds in the presence of additional phosphate, the kleptoplasts are supplied with ample phosphates and nitrogen to maintain photosynthesis longer (Fig. 2). This could also partly explain why the addition of ammonium had a deleterious effect on the photosynthesis of the kleptoplasts (Fig. 4), but not the chloroplasts in the algae (Supplementary Fig. S3); additional ammonium to the already ammonium rich environment of the sea slug cell is too much for the kleptoplasts.

The importance of carbon fixation via kleptoplasts to benefit the energetic needs of the sea slug host has long been contested (Christa et al. 2014), which has led to suggestions that perhaps the importance of kleptoplasts is not only linked to the production of sugars, but also to help in nitrogen cycling (Cartaxana et al. 2021). Indeed, it is even possible that the primary role for the kleptoplasts is to work as “garbage mills” of nitrogenous waste that supply the host with a more diverse repertoire of nitrogen assimilation pathways that are beneficial especially during starvation. However, as sea slugs starved in the presence of an additional nitrogen source tended to fare worse than the control sea slugs (Fig. 4), the nitrogen recycling capacity of kleptoplasts seems to be limited. Regardless, this would also have one very important implication regarding the longevity of the kleptoplasts, as Fd-GOGAT could serve as an alternative electron sink for photosynthesis, but the prominence of this pathway has not been tested in starvation experiments. Interestingly, there are multiple reports showing that photosynthetic electron transfer of PSII remains functional in the sea slugs for weeks and months (Figs. 1, 2, 3) (e.g., Händeler et al. 2009; Cartaxana et al. 2023), but the carbon fixation of the kleptoplasts seems to decrease at a much higher rate (Clark et al. 1981; Marin and Ros, 1989; de Vries et al. 2015). If photosynthetic electron transfer is functional even though the main electron sink, carbon fixation in the CBB cycle is not, where do the electrons go to avoid overreduction of the electron transfer chain and production of deleterious reactive oxygen species? The hypothesis that nitrogen cycling by GS-GOGAT is the main alternative electron sink in starving sea slugs is relatively easy to test, and the results have the potential to answer some fundamental aspects of kleptoplasty in the sea slugs, both in terms of the benefits of kleptoplasts to the animals and how the kleptoplasts are maintained functional. Additional phosphate would also be beneficial, as photosynthetic electron transfer is not expected to proceed long without production of ATP via photophosphorylation, and nitrogen assimilation in the GS-GOGAT pathway is an ATP consuming process. In fact, it was recently suggested that ATP production by the kleptoplasts could be one of the major signals from kleptoplasts to kleptosomes that mark the kleptoplasts as functional or not functional and could determine whether a kleptosome merges with lysosomes to degrade the kleptoplasts (Allard et al. 2025), highlighting the importance of phosphate.

## Conclusions

There are many studies on kleptoplastic sea slugs where the longevity of the kleptoplasts has been assessed by placing the sea slugs away from their food algae to starve. While the effect of multiple different factors to the photosynthetic functionality of the kleptoplasts has been tested, like e.g. temperature (Laetz & Wägele, 2018) and light intensity and fluctuation (Christa et al. 2018; Havurinne & Tyystjärvi, 2020; Laetz et al. 2024), our current study uncovers the hitherto unrealized importance of macronutrients to kleptoplasts. Our results show that if the absolute maximum longevity of the kleptoplasts in the sea slugs is a desired parameter, at least the provision of phosphates should be taken into consideration, as they have a strong positive effect on the longevity of the kleptoplasts. Furthermore, the sea slugs seem to provide the kleptoplasts with phosphate from their own reserves, as well as from the environment to the extent that the photosynthetic activity of the kleptoplasts increases considerably. However, the sea slugs have a limited capacity to repair the kleptoplasts that have been deprived of nitrogen, possibly partially due to the lack of genes required for Chl synthesis. Additional nitrogen had no significant effect on (nitrate) or was even deleterious (ammonium) to kleptoplast longevity, whereas the algae themselves were shown to be able to utilize both forms of nitrogen to completely recover their photosynthetic activity after nitrogen depletion. Nitrogen is undoubtedly important for photosynthesis and kleptoplasts, but our results may indicate that the nitrogenous environment inside the sea slug cells is already supplying the kleptoplasts with plenty of nitrogen, and additional ammonium offsets the nutrient balance of the host and the kleptoplasts. Future research will need to address the importance of the algal GS-GOGAT nitrogen assimilation pathway as an alternative electron sink for the kleptoplasts, both in the presence and absence of phosphate.

## Funding

This work was supported by the European Research Council (ERC) under the European Union’s Horizon 2020 research and innovation programme, grant agreement no. 949880 to S.C. (DOI: 10.3030/949880), and by Fundação para a Ciência e Tecnologia, grants 2020.03278.CEECIND to S.C. (DOI:10.54499/2020.03278.CEECIND/CP1589/CT0012), CEECIND/01434/2018 to P.C. (DOI: 10.54499/CEECIND/01434/2018/CP1559/CT0003), and UID/50006 + LA/P/0094/2020 (DOI:10.54499/LA/P/0094/2020) to CESAM. H.M. received funding from the Finnish Cultural Foundation through the Foundations’ Post Doc Pool.

## Acknowledgements

We thank Maria Inês Silva and Diogo Marçal for technical support in algae and sea slug maintenance. Maria Inês Silva and Axelle Rivoallan are also thanked for their help with the measurements during the starvation experiments.

## Author contributions

V.H.: conceptualization, formal analysis, investigation, methodology, resources, supervision, validation, visualization, writing – original draft. H.M.: funding acquisition, investigation, methodology, writing – review & editing. C.E.: investigation, writing – review & editing. P.C.: funding acquisition, resources, supervision, visualization, writing – review & editing. S.C.: funding acquisition, project administration, resources, supervision, visualization, writing – review & editing.

## Supplementary information

**Supplementary figure S1.**
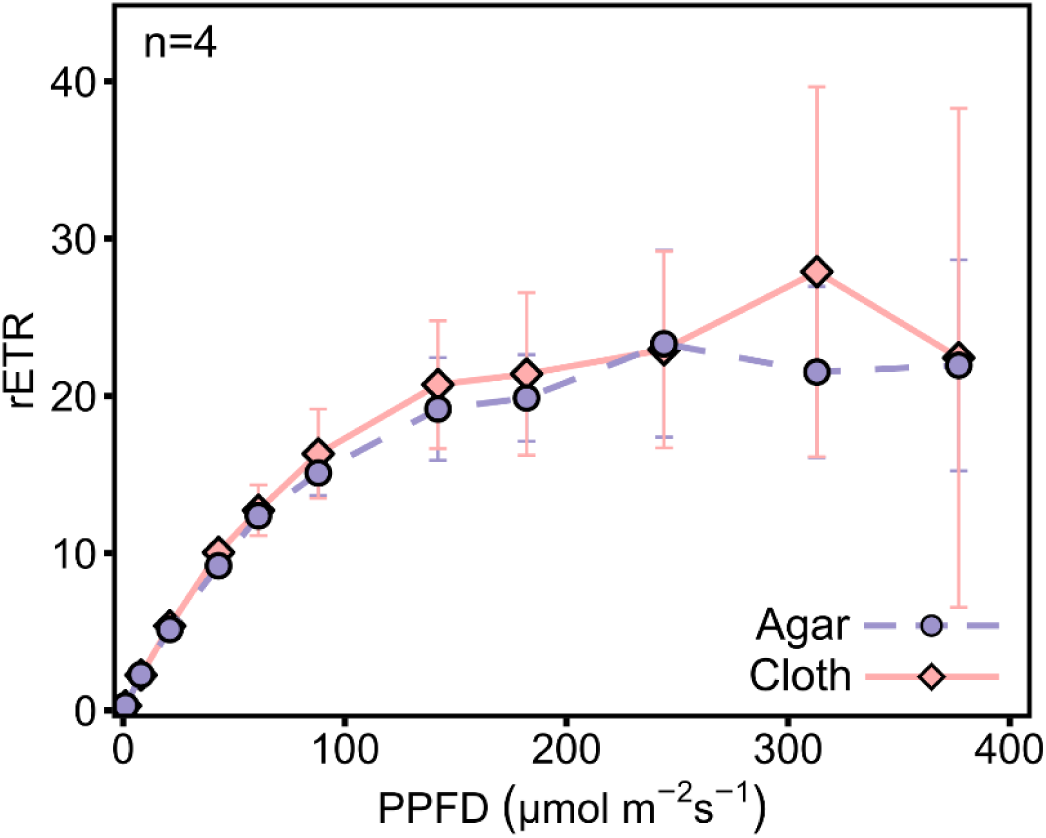
The effect of immobilization method (0.2% agar and cloth) on rETR in the sea slug *Elysia timida*. Relative electron transfer of PSII (rETR) during rapid light curve measurements at the indicated photosynthetic photon flux densities (PPFD) from sea slugs taken from growth conditions and immobilized by embedding them in 0.2% agar or by placing them on a black cloth. The light treatment at each light intensity lasted 60 s before a saturating light pulse was fired to estimate rETR. The data are means from the number of biological replicates shown in the panel. Error bars show standard deviation.

**Supplementary figure S2.**
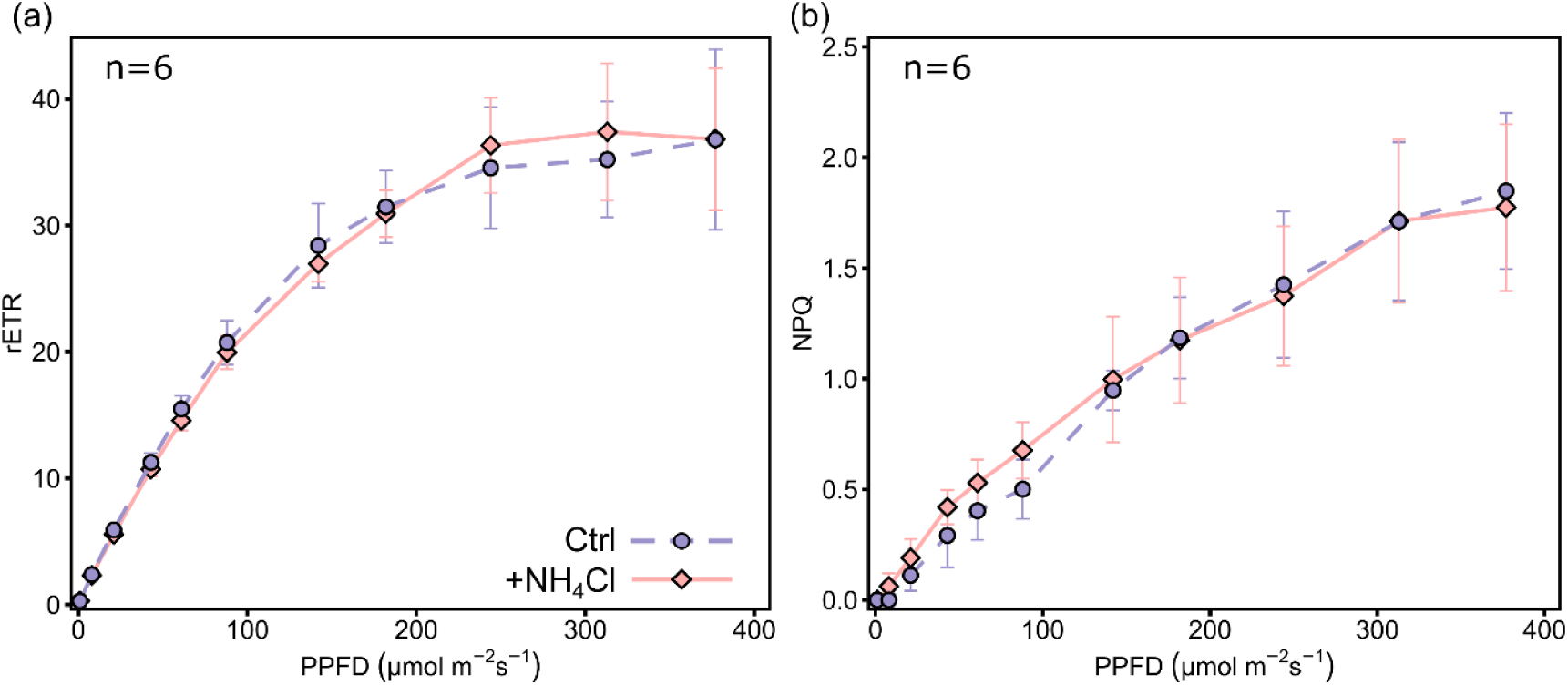
The effect of NH_4_Cl on rETR and NPQ in the sea slug *Elysia timida*. (a) Relative electron transfer of PSII (rETR) during rapid light curve measurements at the indicated photosynthetic photon flux densities (PPFD) from the control and the +NH_4_Cl groups, incubated in regular artificial seawater (ASW) or ASW with 20 μM NH_4_Cl, respectively, for one hour prior to the measurements. (b) The non-photochemical quenching (NPQ) of excitation energy during the rapid light curve, calculated as F_M_/F_M_’−1, where F_M_ and F_M_’ are the maximum fluorescence of PSII at the beginning and during the light treatments, respectively. The first 55 min of the incubation period was done in growth light conditions, while the final 5 min period was in the dark to allow NPQ measurements. The light treatment at each light intensity lasted 60 s before a saturating light pulse was fired to estimate rETR and NPQ. The data are means from the number of biological replicates shown in the panels. Error bars show standard deviation.

**Supplementary figure S3.**
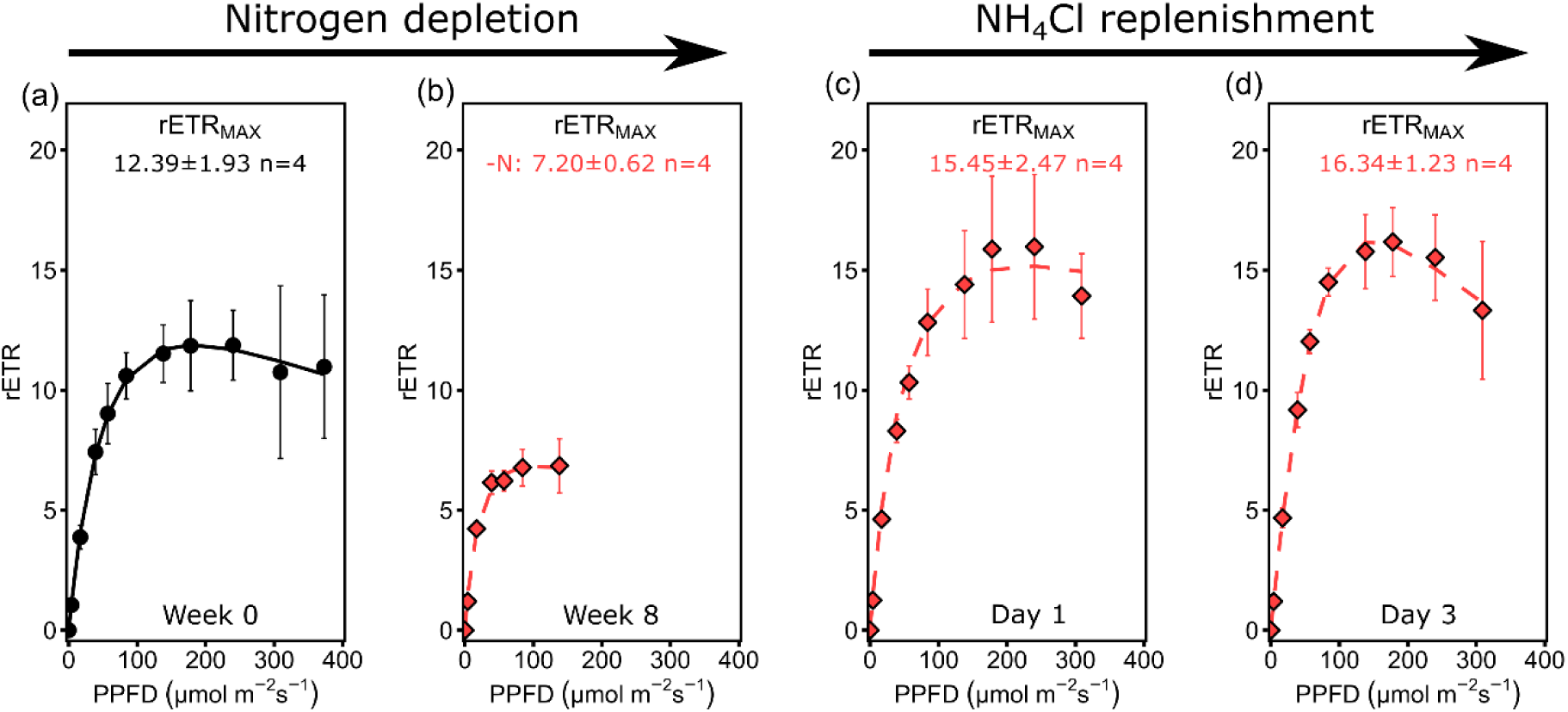
The photosynthetic effect and reversibility of nitrogen depletion with NH_4_Cl in the alga *Acetabularia acetabulum*. (a, b) Rapid light curve (RLC), measuring relative electron transfer of PSII (rETR) at different light intensities (photosynthetic photon flux density; PPFD), from algae at the start of the experiment (a) on week 0 and (b) after 8 weeks without nitrogen. The light treatment at each light intensity lasted 60 s before a saturating light pulse was fired to estimate rETR. The solid and dashed lines show the data modelled according to Eilers and Peeters (1988). Maximum rETR (rETR_MAX_) values derived from the modelled RLCs are shown in each panel. (c, d) RLCs and rETR_MAX_ from nitrogen depleted algae after transfer to vials containing fresh f/2 growth medium with 20 µM NH_4_Cl as a nitrogen source (instead of NaNO_3_) for one and three days. All data are means from the number of biological replicates indicated in the panels. The error bars show standard deviation.

**Supplementary Figure S4.**
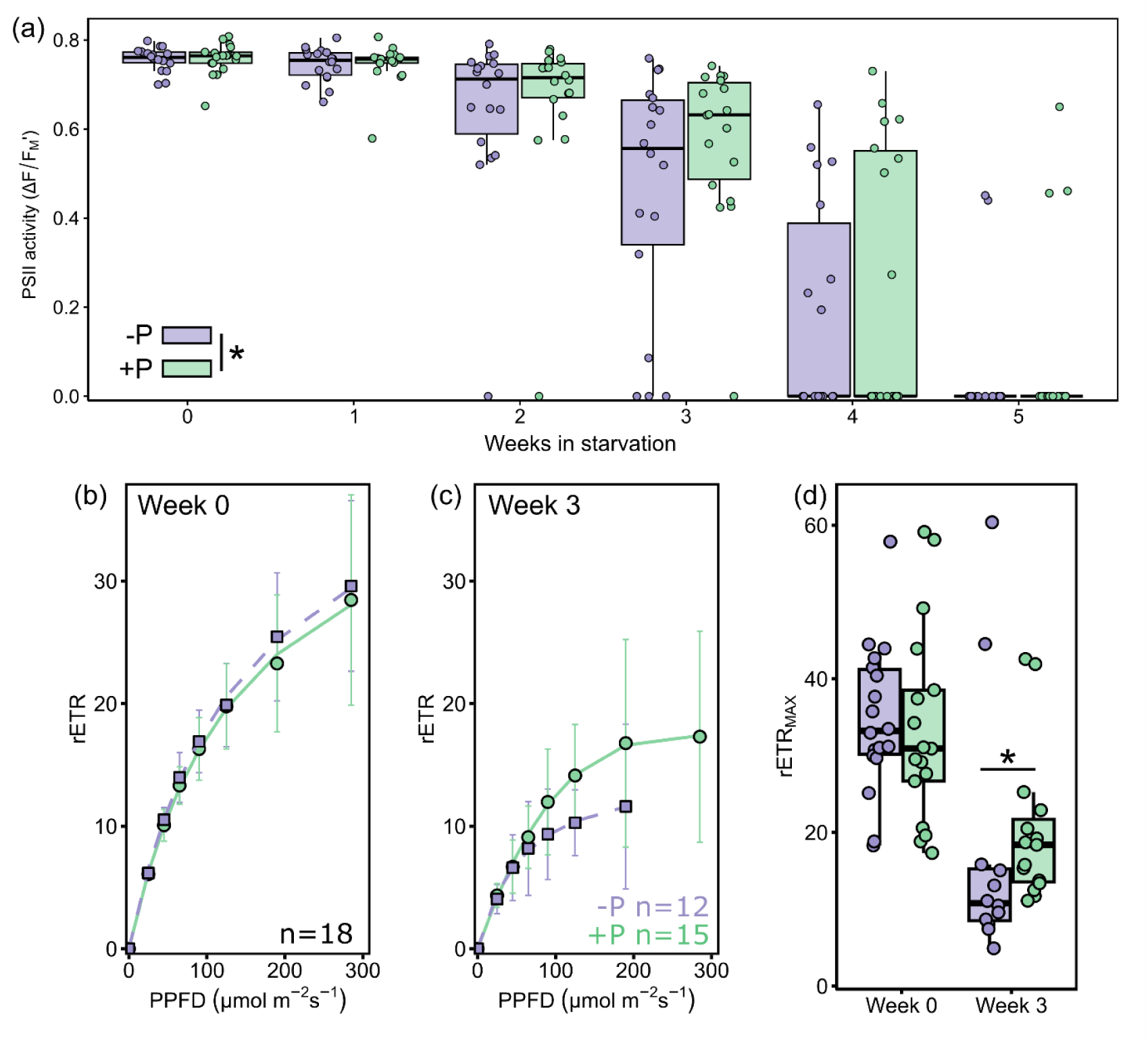
Photosynthetic activity of *Elysia timida* containing phosphate depleted kleptoplasts during long-term starvation in the absence or presence of a phosphate source. (a) The decrease in PSII activity (ΔF/F_M_’) during a 5-week starvation period in the absence (-P; purple) and presence (+P; green) of 36 μM NaH₂PO₄·H₂O. A statistically significant difference in PSII activity caused by the treatment is indicated by * (two-way ART ANOVA, p<0.05). There were no deaths during the five-week starvation period. (b, c) Relative electron transfer of PSII (rETR) during rapid light curve measurements from both groups (b) at the beginning of the experiment on week 0 and (c) on week 3 of starvation, modelled according to Eilers and Peeters (1988) (dashed and solid lines for -P and +P, respectively). Only individuals where a reliable rapid light curve could be measured were used. The light treatment at each light intensity (photosynthetic photon flux density; PPFD) lasted 60 s before a saturating light pulse was fired to estimate rETR. The data are means from the number of biological replicates shown in the panels. Error bars show standard deviation. (d) The rapid light curve parameter rETR_MAX_ (maximum relative electron transfer of PSII) in both treatments at week 0 and week 3 of the starvation. rETR_MAX_ was derived from modelling the data in panels (b, c). ΔF/F_M_’ and RLCs were measured with Junior-PAM (Heinz Walz GmbH) instead of Imaging-PAM used in all other experiments of the study. A significant difference between treatments is indicated by * (two-tailed Wilcoxon rank sum test, p<0.05). The box plots in panels (a, d) show the medians and interquartile ranges, the whiskers indicate non-outlier maxima and minima, and each circle is an individual sea slug. Note that the RLCs on week three could only be reliably measured from 12 individuals of the -P group compared to 15 individuals of the +P group, which suggests even larger differences between the treatments than what is shown in panels c and d.

## Notes

### Competing Interest Statement

The authors have declared no competing interest.

